# The cis-regulatory logic integrating spatial and temporal patterning in the vertebrate neural tube

**DOI:** 10.1101/2024.04.17.589864

**Authors:** Isabel Zhang, Giulia LM Boezio, Jake Cornwall-Scoones, Thomas Frith, Ming Jiang, Michael Howell, Robin Lovell-Badge, Andreas Sagner, James Briscoe, M Joaquina Delás

**Affiliations:** The Francis Crick Institute, London, NW1 1AT, UK; Friedrich-Alexander University Erlangen-Nuremberg, Erlangen, GERMANY

**Author notes:** Correspondence to either or.

## Abstract

The vertebrate neural tube generates a large diversity of molecularly and functionally distinct neurons and glia from a small progenitor pool. While the role of spatial patterning in organising cell fate specification has been extensively studied, temporal patterning, which controls the timing of cell type generation, is equally important. Here we define a global temporal programme operating in progenitors throughout the vertebrate nervous systems that governs cell fate choices by regulating chromatin accessibility. Perturbation of this cis-regulatory programme affects sequential cell fate transitions in neural progenitors and the identity of their progeny. The temporal programme operates in parallel to spatial patterning, ensuring the timely availability of regulatory elements for spatial determinants to direct cell-type specific gene expression. These findings identify a chronotopic spatiotemporal integration strategy in which a global temporal chromatin programme determines the output of a spatial gene regulatory network resulting in the temporally and spatially ordered allocation of cell type identity.

## Introduction

The developing vertebrate neural tube generates a remarkable diversity of neuronal and glial subtypes. This diversity is essential for neural circuit formation and function, yet how it arises during development remains incompletely understood. A gene regulatory network establishes spatial patterns of transcription factor expression in neural progenitors that governs the identity and pattern of neuronal and glial subtype generation (Bella et al., 2021; Cadwell et al., 2019; Frith et al., 2023; Tiberi et al., 2012). But spatial patterning alone fails to explain the full diversity of cell types. Individual progenitor domains produce molecularly distinct neurons and glia in a specific sequential order, markedly expanding neural cell diversity (Delile et al., 2019; Gao et al., 2014; Lim et al., 2018; Lyu et al., 2021; Manno et al., 2021; Mayer et al., 2018; Sagner et al., 2021; Telley et al., 2019; Xu et al., 2014).

The intersection of spatial and temporal identity not only increases molecular diversity but also organises the arrangement of functionally distinct cell types in both vertebrates and invertebrates (Butt et al., 2005; Erclik et al., 2017; Konstantinides et al., 2022; Sen, 2023; Sockanathan and Jessell, 1998; Bayraktar and Doe, 2013). For example, long-range projection neurons in the spinal cord are generated prior to locally connected interneurons but the subtype of projection or local interneuron is determined by the position of generation (Deska-Gauthier et al., 2024; OssewardII et al., 2021). This suggests that spatial and temporal information are integrated within neural progenitors to allocate the spatially and temporally appropriate neuronal subtype. But how temporal patterning is executed in progenitors and how spatial and temporal information are integrated to direct cell fate decisions is unclear.

In the *Drosophila* nervous system, a set of spatial patterning transcription factors (STFs) establish chromatin accessibility configurations in different neuroblasts (Sen et al., 2019; Bayraktar and Doe, 2013; Erclik et al., 2017). These provide the context for cascades of temporal transcription factors (TTFs) to bind chromatin and specify different cell fates (Doe, 2017; Kohwi and Doe, 2013; Sen et al., 2019). This suggests a spatiotopic mechanism for the integration of spatial and temporal information: STFs determine the set of cis-regulatory elements (CREs) accessible in a particular neuroblast thereby governing the binding profile of TTFs and hence the specific gene expression programme activated.

In the vertebrate nervous system, the transition from the production of early-born neurons to later-born neuronal subtypes and glia has provided insight into the temporal programme. Early-born neurons, throughout the nervous system, express TFs of the Onecut family, irrespective of the dorsal ventral progenitor domain from which they are generated. A subsequent wave of neurons express *Pou2f2* and *Zfhx2*–*4*, and then a wave of *Nfia*/*b*/*x* and *NeuroD2* expressing neurons are produced (Clark et al., 2019; Delile et al., 2019; Javed et al., 2020; Manno et al., 2021; Moreau et al., 2021; Sagner et al., 2021). During this final wave, progenitors transition to the production of astrocytes and oligodendrocytes, suggesting that the switch from neurogenesis to gliogenesis is part of a unified temporal patterning programme (Rowitch and Kriegstein, 2010; Sagner et al., 2021).

Temporal changes in the identity of neurons and glia are accompanied by changes in gene expression in neural progenitors. In the vertebrate retina, a conserved transcriptional cascade in progenitor cells controls the specification of differentiated cell types (Cepko, 2014) (Mattar et al., 2015). Similarly, progenitors in the forebrain (Lodato and Arlotta, 2014; Moreau et al., 2021; Telley et al., 2019) and midbrain (Deng et al., 2011; Tiklová et al., 2019) are characterized by a temporal sequence of transcriptional states. Across the neuraxis, *Sox9* and *Nfia*/*Nfib* are sequentially induced in progenitors as they switch from generating early-born to later-born neuronal subtypes and glial cells (Scott et al., 2010; Stolt et al., 2003; Chaudhry et al., 1997; Deneen et al., 2006; Plachez et al., 2008). Sox9 and Nfia/Nfib play a functional role in controlling the transition from neurogenesis to gliogenesis (Deneen et al., 2006; Stolt et al., 2003; Tchieu et al., 2019). However, to what extent the progenitor temporal programme controls the different waves of neurogenesis and which aspects of this regulation are conserved across the central nervous system remains to be determined.

A set of cis-regulatory elements (CREs) responsible for directing gene expression in neural progenitors has been identified (Delás et al., 2023; Kutejova et al., 2016; Nishi et al., 2015; Oosterveen et al., 2012; Peterson et al., 2012). In contrast to *Drosophila* (Sen et al., 2019; Xu et al., 2024), most neural progenitors share a common chromatin landscape, with minimal variation in the accessibility of CREs across progenitor types, at the early stages of neural tube patterning, despite the significant functional and gene expression differences. The similarity in the chromatin landscape between spatially distinct neural progenitors appears to rule out a spatiotopic mechanism. An alternative, therefore, is a chronotopic strategy in which TTFs modify chromatin accessibility in progenitors thereby altering the output of STFs and consequently the cell types specified over time.

To test this possibility and to define the gene regulatory logic integrating spatial and temporal patterning in the vertebrate neural tube we used chromatin accessibility assays and functional perturbations. The analysis revealed a temporal patterning cascade comprising a set of TTFs that are sequentially induced, independently from spatial inputs, and act to modify chromatin accessibility and alter gene expression over time. Thus, by contrast to *Drosophila*, in the vertebrate neural tube, TTFs establish the chromatin context for spatial determinants to specify cell type identity. This defines a novel mechanism for generating and organising cell fate diversity in the developing nervous system.

## Results

### A temporal programme of chromatin accessibility in neural tube progenitors

The spatially restricted expression of a set of transcription factors (TFs) partitions neural progenitors of the developing neural tube into discrete domains of progenitors arrayed along the dorsal ventral axis (Fig 1A, (Dessaud et al., 2008)). A distinct set of temporal transcription factors (TTFs) defines sequential phases of neural progenitor development (Sagner et al., 2021). These are expressed in neural progenitors along the entire dorsal ventral axis (Fig 1A). To understand how neural progenitors integrate their dorsoventral spatial identity with the temporal progression of cell fate changes, we made use of an *in vitro* cellular model based on the directed differentiation of embryonic stem cells (ESCs) (Delás et al., 2023; Gouti et al., 2014) (Fig 1B). We assayed *in vitro* generated ventral neural progenitors (NPs) from day 5 until day 11 of differentiation. Gene expression recapitulated the temporal hallmarks expected from *in vivo*, including sequential onset of expression of the transcription factors *Sox9*, *Nfia* and *Nfib* (Fig S1A), and the later appearance of glia markers *Slc1a3* (*Glast)* and *Gfap* (Fig S1B). Expression of NFIA at the protein level was detected from day 9, reaching 90% of SOX2+ NPs by day 11 (Fig S1E).

**Figure 1:**
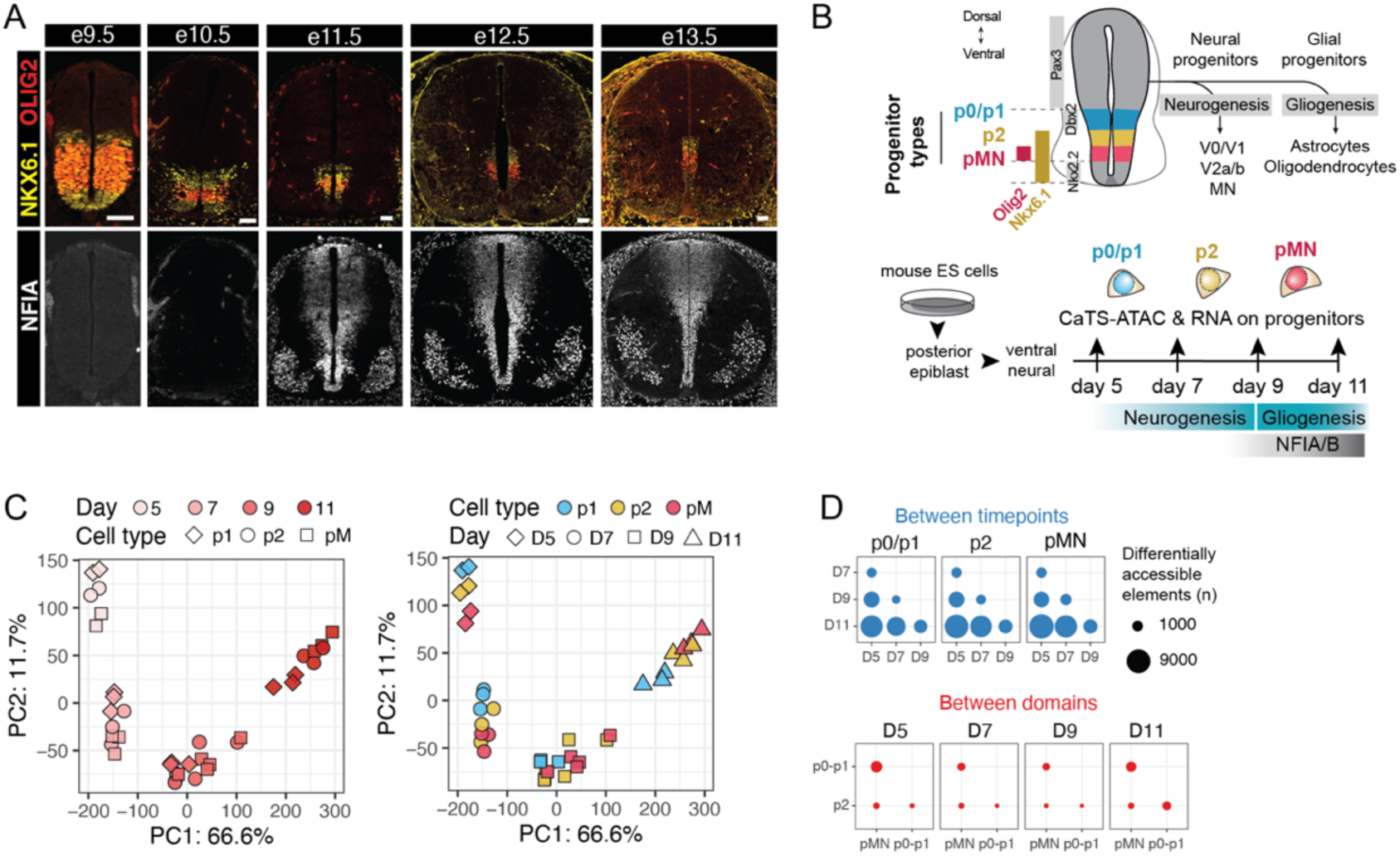
A global chromatin temporal programme in spinal cord progenitors. (A) Expression of spatial transcription factors NKX6-1 and OLIG2 across all timepoints from mouse embryonic day 9.5 to 13.5. The temporal transcription factor NFIA is first detected at e11.5. Scale bar 50 µm. (B) Diagram of the neural tube progenitor subtypes characterised in this study and the differentiated cell types they produce (top). Diagram of the in vitro system and the time points assayed (bottom). (C) Principal component analysis (PCA) of CaTS-ATAC for all timepoints and cell types. The main changes are seen between timepoints (samples coloured by day, left), for all cell types analysed (coloured by cell type, right). (D) Quantification of the number of differentially accessible regions between timepoints for each cell type (top) and between cell types for all pairwise cell type comparisons at each time point (bottom) highlights the large number of changes detected over time.

We focused on progenitor cells with three spatial identities, p0-p1, p2 and pMN. The three progenitor domains generate a variety of temporally distinct neurons (motor neurons (MNs) and interneurons) and later glial cell types (oligodendrocytes and astrocytes) (Briscoe et al., 2000; Deneen et al., 2006; Hayashi et al., 2018; Hochstim et al., 2008; Lu et al., 2000; Novitch et al., 2001; Worthy et al., 2023; Zhou et al., 2000). This suggests a temporal programme must intersect with the spatial identity of progenitors to determine the fate of progeny. To investigate this, we first performed cell-type-specific ATAC and RNA-seq using our previously developed method, CaTS-ATAC (Delás et al., 2023), separating progenitor cells of either p0-1, p2 or pMN identity at days 5, 7, 9, and 11. These three progenitors can be generated in the same signalling conditions and isolated based on a combinatorial code of transcription factor expression: all express SOX2, but pMN cells express OLIG2 and NKX6-1; p2 cells express NKX6-1 only; and p0-p1 do not express either of these TFs (Fig S1C,D). In these conditions, there is little or no induction of more dorsal (PAX3) or more ventral (NKX2-2) progenitors (Fig S1F,G).

Consistent with our previous study, the chromatin accessibility was similar in the three progenitor domains at day 5 (Fig 1C) (Delás et al., 2023). Moreover, the three progenitor types had similar chromatin accessibility profiles at each time point. However, the set of accessible regions differed between time points (Fig 1C, D). A clear, strong, global temporal signature of chromatin accessibility dominates the chromatin landscape (Fig 2A). Consequently, the number of differentially accessible elements in the same cell type at different timepoints is far greater than the differences in chromatin accessibility between cell types at the same time point (Fig 1D). Taken together, the data show a strong temporal programme of chromatin accessibility that is shared between all three progenitor domains examined.

### A global temporal chromatin programme operates in vivo throughout the neuraxis

We next asked whether the temporal differences in chromatin accessibility observed *in vitro* were also present *in vivo* and extend beyond the spinal cord to the rest of the developing nervous system. Analysis of *in vivo* spinal cord scATACseq (Shu et al., 2022) showed similar temporal chromatin accessibility changes in neural progenitors as observed *in vitro*. Early chromatin elements were accessible at e9.5 *in vivo*, while late chromatin elements opened at later timepoints (Fig 2A, B). The presence of progenitors from across the dorsoventral axis at each timepoint highlighted that the temporal chromatin programme operates across spatial domains *in vivo*. This indicates that *in vitro* differentiation accurately recapitulates *in vivo* developmental transitions and suggests a consistent progression of temporal transitions in chromatin accessibility between *in vitro* and *in vivo* conditions.

**Figure 2:**
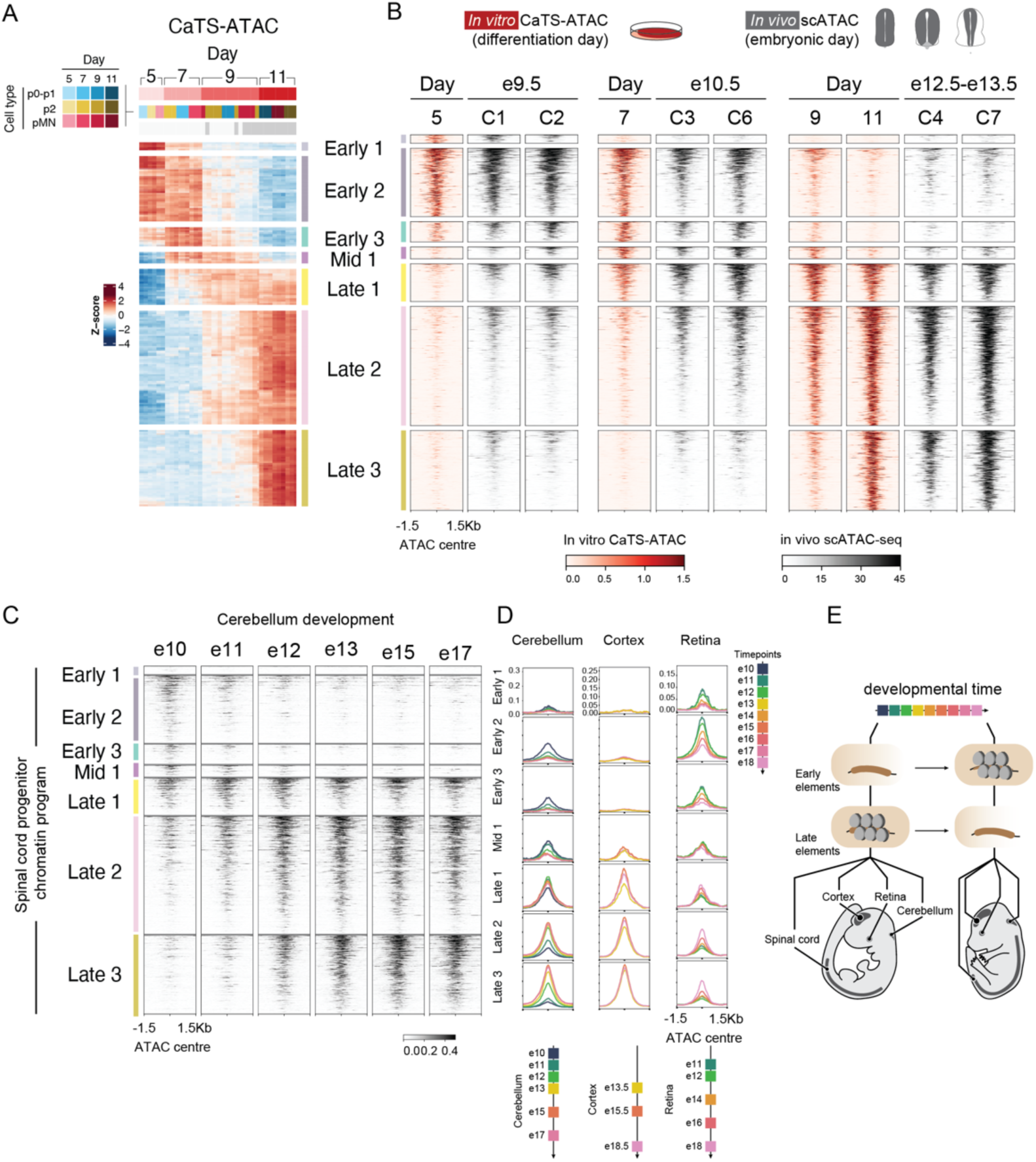
The chromatin temporal programme is conserved across the central nervous system. (A) Heatmap of high confidence differentially accessible elements over time, shared in all cell types. (B) Comparison of the temporal elements between in vitro differentiation (this study) and in vivo scATAC seq (Shu et al., 2022). (C) The same elements display the same temporal ordering of opening and closing in cells from the cerebellum (Sarropoulos et al., 2021). (D) Equivalent accessiblity dyanmics for these elements was evident in the developing cortex (Bella et al., 2021) and retina (Lyu et al., 2021). (E) Diagram highlighting how accessibility at a set of chromatin regions changes over time across the central nervous system.

Previous studies of more rostral regions of the developing nervous system have identified chromatin elements differentially accessible in early or late progenitors (Lyu et a., 2021; Sarropoulos et al., 2021). We therefore examined whether the chromatin programme identified in the spinal cord is conserved across the rostral-caudal axis. In the cerebellum (Sarropoulos et al., 2021), early temporal elements were open at e10-e11 but closed at later timepoints, while late temporal elements progressively opened between e11 and e13 (Fig 2C, D). Similar results were observed in the retina (Lyu et al., 2021) (Fig 2D). Data from cortical cells, starting at e13.5 (Bella et al., 2021), showed late elements were open (Fig 2D). Moreover, region-specific re-analysis of an organogenesis atlas (Sun et al., 2024) showed that early elements were open at e10.5-11.5 in di-telencephalon, mesencephalon, hindbrain, and spinal cord cells (Fig S2A) before closing and late elements opening.

These findings demonstrate that the temporal chromatin programme identified in the spinal cord operates globally across central nervous system regions. This raises the possibility that the same transcription factors act via identical regulatory regions, control chromatin accessiblity across the neuraxis to coordinate temporal progression (Fig 2E). Moreover, previous comparative analyses (Lyu et al., 2021; Sarropoulos et al., 2021; Yuan et al., 2022) suggest that this regulation is likely to be conserved across species, including humans.

### A screen identifies regulators of progenitor temporal progression

We hypothesized that transcriptional factors with temporally changing expression could drive the changes in chromatin accessibility, NPs of different spatial domains share a temporal transcriptional programme that is conserved across the rostral caudal (Sagner et al., 2021). This programme includes around 30 transcription factors, the expression of which defines sequential developmental phases(Sagner et al., 2021). Our CaTS-RNA data recapitulated this sequence: early expression of *Lin28a*/*Lin28b* and *Nr6a1* was followed by expression of *Npas3* and *Sox9* and then later expression of Nfia/Nfib (Table S3). Gene ontology analysis highlighted differential enrichment of biological processes, with gene clusters expressed early being enriched for developmental terms, whereas late gene clusters were enriched for neuronal processes (axonogenesis, synapse organisation) (Fig S1H). This is consistent with the temporal transcriptional programme transition described for apical progenitors (Telley et al., 2019).

To explore the involvement of temporal transcriptional factors in the changes in chromatin accessibility, we examined differences in motif usage between different times using footprinting to identify motifs likely to be bound by proteins. We used TOBIAS (Bentsen et al., 2020) and grouped footprint-containing motifs into archetypes based on PWM clustering (Vierstra et al., 2020). The analysis revealed several motifs that were predicted to be bound differentially between timepoints, and correlated with changes in gene expression for TFs that bind to these motifs. For example, the expression of *Rfx4* correlated with footprints on an RFX motif (Fig 3A,B). At later time points, we see the expected expression and footprint for NFIA TFs. *Sox9* is known to be upregulated at an intermediate stage of neural progenitor development (Stolt et al., 2003) and indeed its expression correlated with the presence of a SOX footprint. However, several other SOX TFs are expressed across this temporal transition and could contribute to the observed footprint (Fig S1I). We also identified the AP1 motif as differentially bound and several TFs that bind this motif are differentially expressed over time (*Atf3*, *Fos*, *Junb*). These could individually or in combination explain the observed footprint (Fig 3A,B).

**Figure 3:**
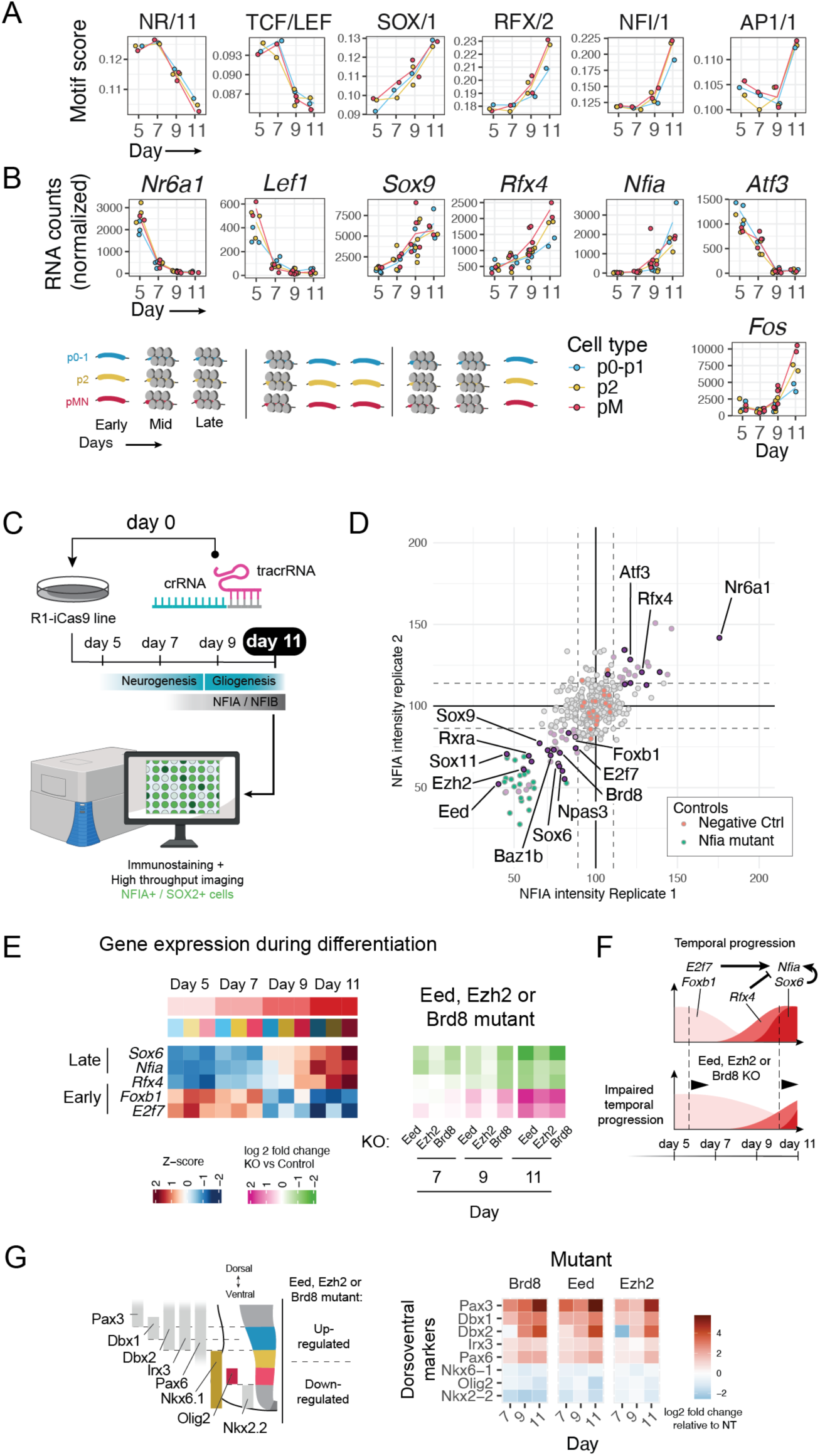
A CRISPR screen identifies regulators of the global temporal program. (A) Motifs with differential footprints over the temporal progression. (B) Expression levels of TFs with similar temporal dynamics that could bind those motifs. Both dynamic footprints and gene expression occur in all cell types in a coordinated manner. (C) Diagram of the CRISPR screen set up and readout. (D) Results across two biological replicates with controls and some hits highlighted. (E) Average expression (CaTS-RNA) of the indicated genes across cell types and timepoints (left). Fold change expression upon mutation of *Eed, Ezh2* or *Brd8* for the same genes at three timepoints. (F) Diagram summarising the effects of *Eed, Ezh2* and *Brd8* on the temporal program. Late genes are downregulated and earlier genes upregulated, consistent with a delayed temporal progression. (G) Diagram of the expression of spatial transcription factors (left). Dorsal genes are upregulated when *Ezh2, Eed* or *Brd8* are mutated compared to non-targeting controls. Nr6a1 promotes the early progenitor programme and inhibits temporal progression.

These analyses highlighted candidate TFs that may drive the temporal programme in NPs. However, a single footprint can result from the effects of multiple TFs and not all TFs produce identifiable footprints. Moreover, many components of chromatin-modifying complexes are differentially expressed over time but are not associated with specific genomic motifs. To define drivers of the temporal programme, we performed an arrayed CRISPR mutation screen. We selected ∼300 differentially expressed genes during the temporal progression, including TFs, chromatin modifiers and remodellers and members of signaling pathways. We used high-content imaging and quantified NPs (SOX2+) at day 11 of differentiation expressing the late NP marker NFIA to identify to regulators of temporal progression (Fig 3C).

The screen identified several positive regulators of NFIA, targeting of which led to a depletion in NFIA levels, similar to targeting *Nfia* itself (Fig 3D, Fig S3A). This included *Sox9*, which is known to be required for the temporal programme and the expression of NFIA *in vivo* (Kang et al., 2012). In addition, two subunits of PRC2 (Eed and Ezh2) and two members of chromatin remodelling complexes (Baz1b, Brd8) were identified to be involved in NFIA induction. Conversely, targeting of *Rfx4* and *Nr6a1*, both of which are expressed in NPs earlier than NFIA although starting at different timepoints, resulted in increased NFIA expression (Fig 3D). This suggests that expression of these factors might normally delay NFIA induction.

We first investigated the effects of *Eed* and *Ezh2*, which are members of the polycomb complex, and *Brd8*. To distinguish whether the reduction in NFIA expression observed in the primary screen was due to lower levels of NFIA expression or a lower proportion of cells expressing NFIA, we performed intracellular flow cytometry for NFIA and SOX2. This revealed that mutation of either *Eed*, *Ezh2* or *Brd8* resulted in lower proportion of NFIA+ progenitors, from the onset of NFIA expression at day 9 (Fig S3B). Because the polycomb complex and chromatin remodellers regulate many aspects of cell identity, we performed RNAseq. In agreement with *Eed*, *Ezh2* and *Brd8* promoting NFIA expression, NFIA expression was reduced at days 9 and 11 after targeting *Eed*, *Ezh2* or *Brd8*, compared to control. Consistent with an overall disruption to the temporal programme, genes normally expressed during the early phases (e.g., *Foxb1*) were upregulated at later times and genes normally expressed at late times (e.g., *Rfx4*, *Sox6*) were downregulated (Fig 3E, S2C). These included several of the TFs identified as hits in the screen (Fig 3E). These results suggest a role for the polycomb complex and chromatin remodellers in the progression of the temporal programme in the neural tube (Fig 3F). This is consistent with the extension of the early temporal competence window in *Drosophila* neuroblasts lacking PRC1 or PRC2 (Lucas et al., 2021) although results in mammalian cortex (Amberg et al., 2022; Telley et al., 2019) highlights the complexity of regulating the temporal transitions.

However, mutation of *Eed*, *Ezh2* or *Brd8* also altered expression of dorsoventral identity genes, increasing expression of dorsal markers such as *Pax3*, *Dbx1* and *Dbx2* (Fig 3G), while the expression of the expected ventral markers, such as OLIG2, were abrogated (Fig 3G, S2D). These data are consistent with polycomb group proteins and Brd8 concomitantly affecting the temporal progression and spatial identity of neural progenitors.

The primary screen identified orphan nuclear receptor Nr6a1 as a strong negative regulator of NFIA. Both in vivo and our CaTS-RNA data show *Nr6a1* is expressed at early timepoints and sharply dowregulated by day 7 of differentiation (Fig 2A, 3A, (Sagner et al., 2021)). We therefore wanted to test whether *Nr6a1* played a functional role in the control of the temporal dynamics that we have observed. Targeting *Nr6a1* by CRISPR guide transfection resulted in a higher proportion of NFIA-positive progenitors as early as day 9, confirming *Nr6a1* as a negative regulator of NFIA (Fig 4B). Consistent with a disruption of the global programme, several genes usually induced at day 9 or later (Fig 4D) were upregulated as early as day 7 (Fig 4C, Fig S4A). These included genes known to be part of the gliogenic switch or implicated in gliogenesis such as *Nfib* itself, *Sox10*, *Aldh1l1*, *Bcl11a*, *Bcan* and *Slc1a3* (also known as *Glast*) (Fig 4C). This supports a role of *Nr6a1* as a negative regulator of the temporal programme.

**Figure 4:**
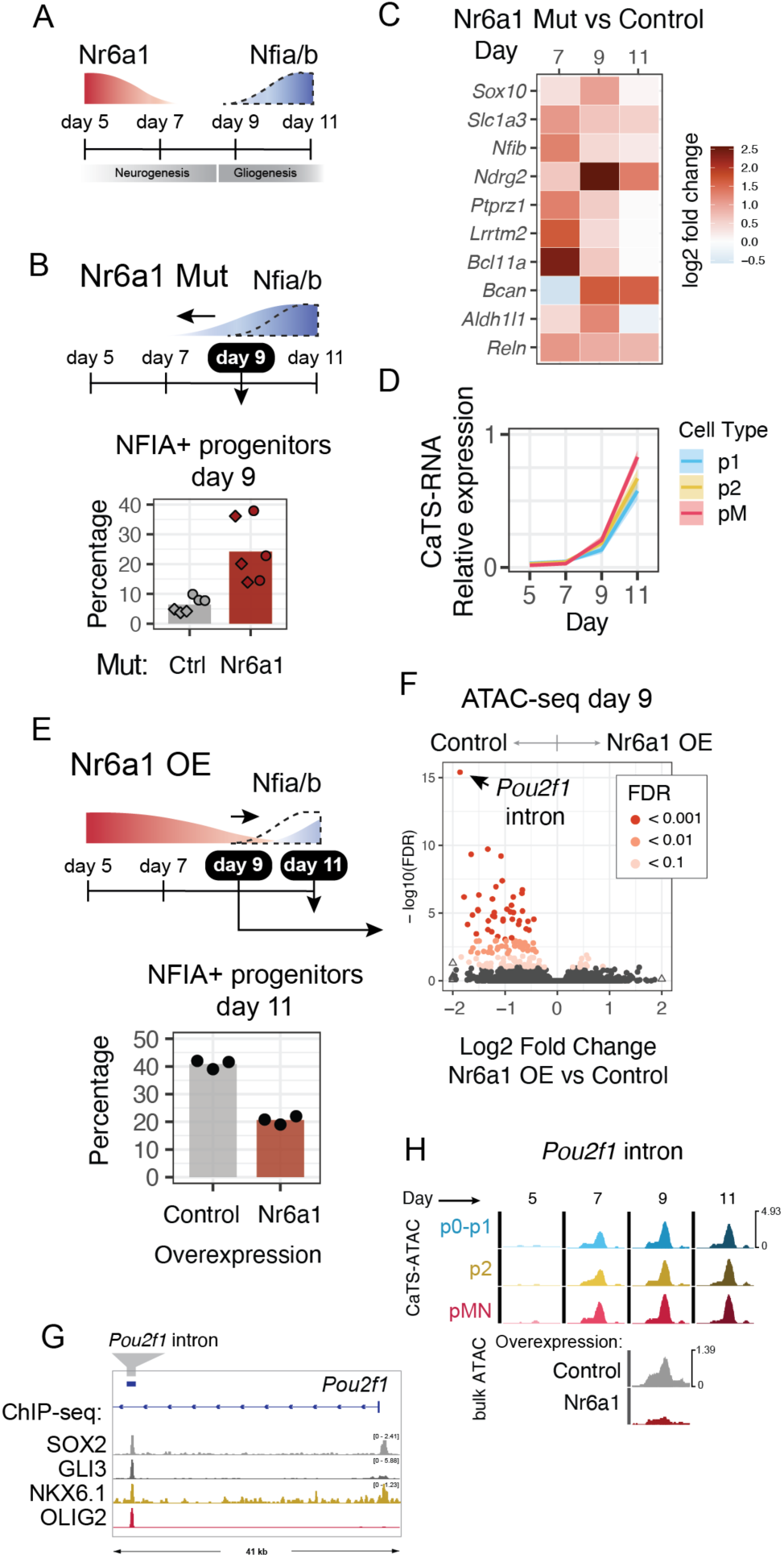
Nr6a1 is a regulator of the temporal program. (A) Diagram of the temporal expression of Nr6a1 and Nfia/Nfib. (B) Nr6a1 KO results in a higher proportion and earlier generation of NFIA+ progenitor cells. (C)Fold change in Nr6a1 KO versus non-targeting shows early upregulation of these genes. (D)Average expression of genes in (C) would usually start at day 9 or 11. (F) Nr6a1 overexpression by lentiviral transduction at day 5 results in reduced proportion of NFIA+ progenitors are day 11. (G) ATAC-seq at day 9 shows that elements that would have usually opened by day 7 or 9 (Fig S4B) open in control but not in Nr6a1 overexpression. (H) Image from the genome browser of the Pou2f1 locus showing an element bound by SOX2, GLI and STF NKX6-1 and OLIG2. (I) CaTS-ATAC of the element from (G) shows the opening of the element in all cell types by day 7 (top). The element opens in control but fails to open in Nr6a1 overexpression (bottom).

We tested whether the reverse, overexpression of *Nr6a1*, would delay temporal progression. To prolong the expression of *Nr6a1* beyond its endogenous window, we used lentivirus expression to transduce differentiating NPs at day 5. Flow cytometry analysis at day 11 showed that the proportion of NFIA positive progenitors was reduced by 50% in Nr6a1 overexpression versus control (Fig 4E). This is consistent with Nr6a1 maintaining cells in an earlier temporal state and acting as a negative regulator of temporal progression.

We reasoned if *Nr6a1* is controlling the temporal programme via changes in chromatin accessibility, we would expect to see changes in accessibility upon *Nr6a1* overexpression. Consistent with this, ATAC-seq at day 9 identified elements that specifically opened in control, but not in cells overexpressing *Nr6a1* (Fig 4F). These elements correspond to dynamic sites that are usually opened by day 7 or 9 in all cell types (Fig S4B). This included an element near *Pou2f1* (Fig 4G, H). Pou2f1 is differentially expressed and has been reported to control temporal progression in *Drosophila* neuroblasts and vertebrate retina (Doe, 2017; Javed et al., 2020).

Together, these data support a role for *Nr6a1* in promoting the early chromatin programme and antagonising the temporal progression of neural progenitors.

### The global chromatin temporal programme directs position-specific cell type gene expression

We next asked how differential gene expression between spatial domains over the temporal progression was controlled, given how all spatial domains shared a global temporal chromatin programme. Our previous work demonstrated that differential binding of cell type specific TFs at commonly accessible elements controls cell type specific gene expression in early NPs. This also appears to be the case at later timepoints. For example, chromatin accessibility associated with *Pax6* (Fig 5A) or *Dbx1* (Fig S5A) is similar in p0-p1, p2 and pMN progenitors at day 5, 7, 9 and 11, despite these genes having distinct spatial expression patterns. A regulatory element documented to control *Pax6* in NPs (Oosterveen et al., 2012) is accessible in all cell types and remains accessible over time. The element is bound by NKX6.1 and OLIG2, which are themselves differentially spatially expressed transcriptional repressors. Thus, NKX6.1 binding in p2 and/or pMN, and OLIG2 binding in pMN can repress the Pax6 expression in these cell types.

**Figure 5:**
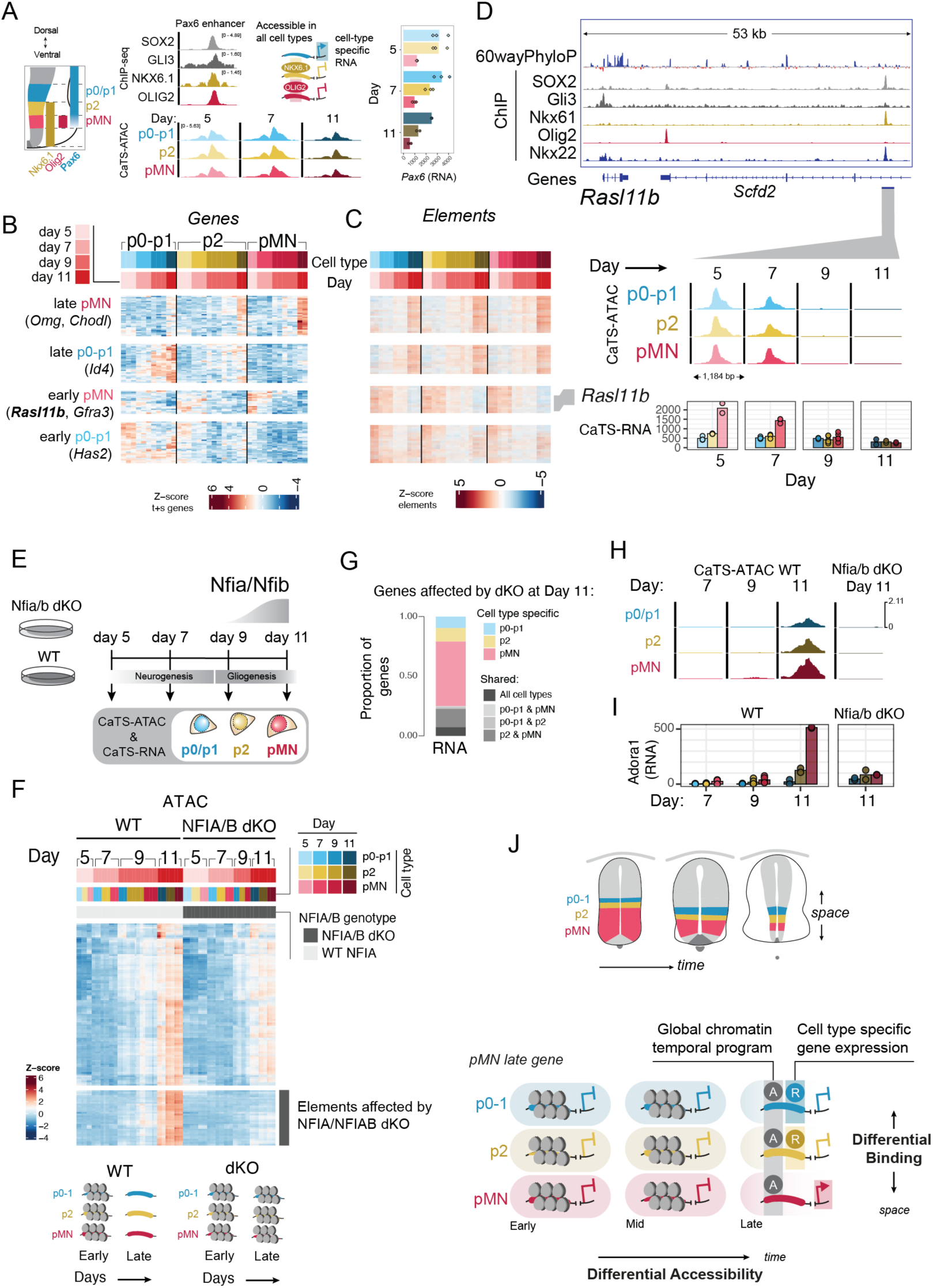
The temporal chromatin programme controls dynamic, cell-type specific gene expression. (A) Spatial Transcription Factor PAX6 is differentially expressed between p0-p1, p2 and pMN (diagram: left, CaTS-RNA: right). A published enhancer element for *Pax6* is bound by STF NKX6.1 and OLIG2, as well as positive input SOX2 and the Shh effector GLI. The element is open across all cell types and all timepoints. (B) Genes dynamically expressed in time and in a cell type specific manner (for full set, see Fig S5B,C) (C) Elements associated with genes in (B), show a temporal dynamic accessibility, but always shared across all cell types. (D) Example element associated with *Rasl11b* is bound by STF NKX6-1 (top) and accessible at days 5 and 7 but not later timepoints*. Rasl11b* is differentially expressed between cell types. (E) Diagram of the *Nfia/Nfib* double KO profiling by CaTS-ATAC and RNA. (F) A subset of elements that should open at later timepoints fails to open in *Nfia/Nfib* dKO. Affected elements are shared for all cell types. (G) Breakdown of genes differentially expressed between control and dKO at day 11 showing proportion of shared versus cell type specific highlights that genes are primarily affected only in one cell type. (H) Example element associated with *Adora1*, which opens at day 11 but fails to open in the dKO in all cell types. (I) *Adora1* is a cell type specific genes (pMN) expressed at day 11. The dKO fails to express this gene in pMN (cell type specific gene effect). (J) Diagram explaining how the temporal and spatial axis integrate time and space via two distinct chromatin strategies.

We reasoned that differential binding within the temporally available elements could explain the integration of the temporal and spatial patterning programmes. We identified groups of genes that were differentially expressed between cell types and between time points (Fig S5B, Table S4). For example, chromatin regions associated with genes expressed at late times but only in pMN (e.g., *Omg*) or only in p0-p1 (e.g., Id4) (Fig 5B) become accessible only at late timepoints, following the genes’ expression dynamics. However, these regions were accessible in all cell types and not exclusively in the cell type in which the associated gene was expressed (Fig 5C, Fig S5C). This pattern is true for genes displaying other temporal dynamics, such as *Rasl11b*, which is differentially expressed in pMN at early timepoints but not at later times. Its gene expression profile correlated with chromatin accessibility at a nearby element, which is accessible in early p0-p1, and p2 progenitors as well as pMN cells. This element is bound by cell type specific TFs, such as NKX6.1 (Fig 5D), which might confer the spatial specificity to *Rasl11b* from shared chromatin accessibility. This supports a model in which the temporal chromatin programme directs the output of the spatial gene regulatory network to achieve dynamic and cell type specific gene expression.

To test this model, we perturbed the well-studied regulators of the temporal transition, NFIA and NFIB. *Nfia* and *Nfib* are dynamically regulated over time, with onset at day 9 and peaking at day 11 which is consistent with the appearance of their footprints (Fig 3A). As expected, cells lacking NFIA/NFIB had defects in generating glial cells durin in vitro differentiation (Fig S5D). If NFIA/NFIB regulate the temporal programme, we would predict shared chromatin accessibility changes across all domains. Additionally, if these temporal elements control cell type specific genes, we predicted that depletion of NFIA/B would show cell type specific defects in gene expression.

We assayed *Nfia*/*Nfib* double knockout (dKO) cell lines across the time course (Fig 5E). This revealed a loss of chromatin accessibility in a substantial number of regions that usually become accessible at day 11. Moreover, most of the affected accessible elements were shared between progenitor cell types (Fig 5F). This is consistent with a role for NFIA/NFIB in the temporal programme. By contrast, the gene expression changes that resulted from loss of NFIA/B, were mostly cell type specific (Fig 5G). This is also highlighted by examination of specific element-gene pairs. For example, the predicted element associated with *Adora1* gains accessibility at day 11 in all cell types in the wild-type but fails to open in Nfia/Nfib dKO cells (Fig 5H). *Adora1* is a cell type specific gene expressed in pMN at day 11 but fails to be expressed in dKO cells (Fig 5I). This indicates that this differentially expressed gene is controlled by the shared chromatin temporal programme.

Overall, the data support a model in which the temporal chromatin programme is encoded by differential accessibility of elements over time, with specific sets of elements available at each time window. Within these sets of elements, differential binding of TFs converts the shared accessibility to cell type-specific gene expression. This highlights how the two patterning axes, time and space, are encoded by different chromatin strategies, differential accessibility and differential binding (Fig 5J). This suggests a logic to the integration of the regulatory information.

### The progenitor temporal programme controls progeny cell type identity

Previous work has identified that neurons born at different times express different temporal TFs regardless of spatial identity (Delile et al., 2019; Sagner et al., 2021): early-born neurons express *Onecut*; intermediate-born neurons express *Zfhx3*; and later-born neurons express *Nfia*/*Nfib* (Fig 6A). To test whether the temporal programme in progenitors determines the temporal identity of differentiated cells, we examined the effect of perturbing the temporal programme in progenitors.

**Figure 6:**
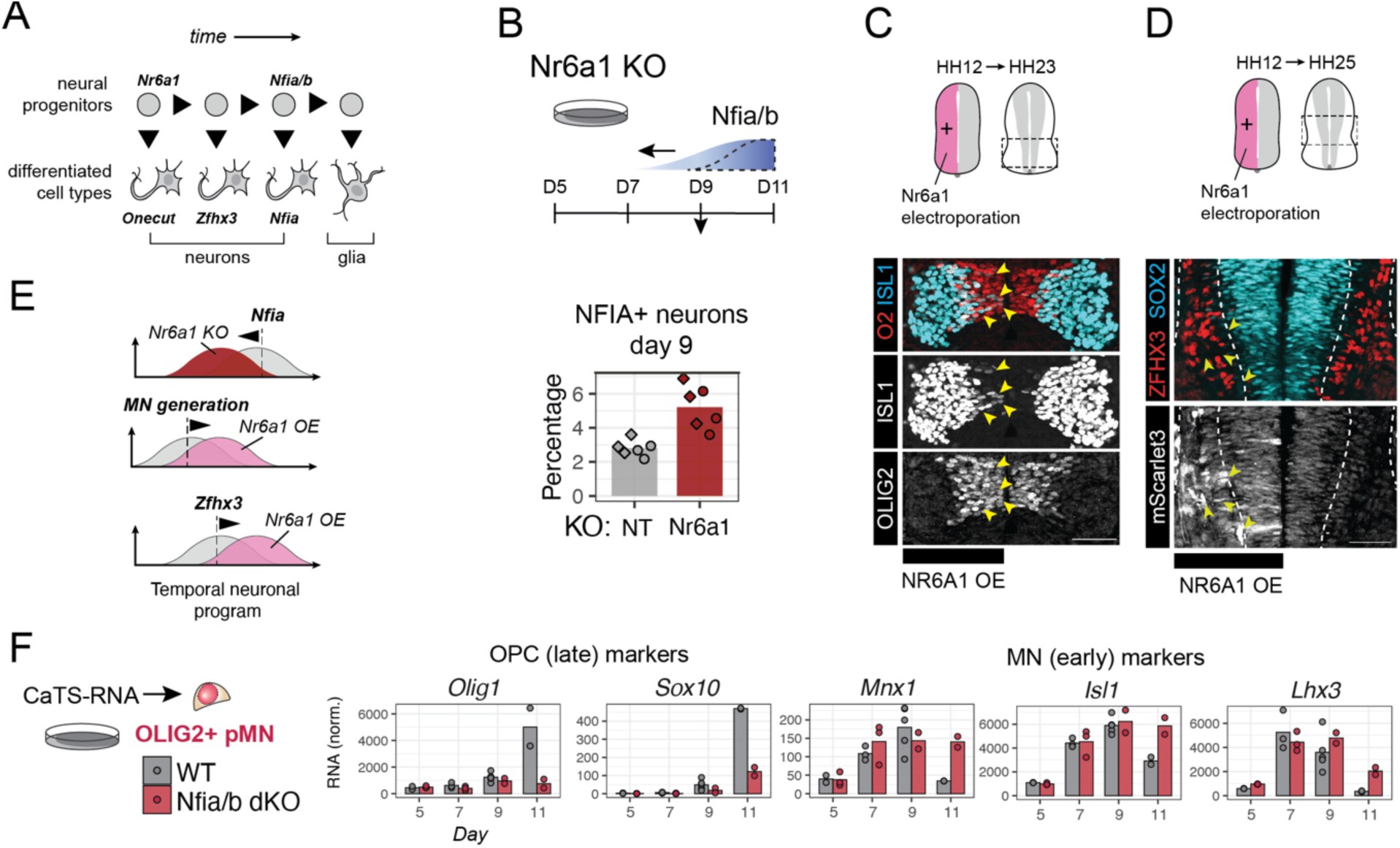
The progenitor programme controls differentiated cell type outcomes. (A) Diagram of the neuronal temporal programme (Delile et al., 2019; Sagner et al., 2021). (B) Nr6a1 KO results in a higher proportion and earlier generation of NFIA+ neurons. (C) Spinal cord from HH23 chicken shows motoneurongenesis still ongoing (ISL1+ OLIG2+ cells) in the side of Nr6a1 electroporation, performed at HH12. (D) Spinal cord from HH25 chicken shows reduced expression of intermediate neuronal marker ZFHX3 in the side of Nr6a1 electroporation. (E) Diagram summarising the effects seen at the neuronal level, with *Nr6a1* KO accelerating late neurons, whereas *Nr6a1* over expression prolongs generation of motor neurons (early) and delays onset of ZFHX3 (intermediate timepoint). (F) Known markers of oligodendrocyte progenitor cells *Olig1* and *Sox10* are not upregulated to the same level as control in cells lacking Nfia/Nfib. Conversely, expression of earlier motor neuron markers is sustained in the dKO (e.g, *Isl1, Mnx1*). Scale bar 50 µm.

Further to the effects seen in progenitors (Fig 4B), *Nr6a1* mutation results in an earlier (day 9) and increased production of NFIA+ (late-born) neurons (Fig 6B). To test the effects on the neuronal programme, we electroporated *Nr6a1* in chick embryos at HH12 (before the onset of neurogenesis), targeting NPs. Consistent with *Nr6a1* promoting the earlier programme and delaying temporal progression, at HH23-24 MNs continued to be generated from *Nr6a1* overexpressed (OE+) pMN cells. This is 12h after the usual cessation of MN generation (Hollyday and Hamburger, 1977). This is evident by the presence of ISL1-expressing cells within the ventricular zone, close to the apical surface of the neuroepithelium, in NPs co-expressing OLIG2 in the electroporated side of the chick neural tube (Fig 6C). Moreover, assaying embryos at HH25, the timepoint when ZFHX3+ neurons (intermediate-born) are first detected, revealed a decrease in ZFHX3+ neurons in areas of *Nr6a1* electroporation, compared to the contralateral side (Fig 6D). This is consistent with *Nr6a1* promoting the earlier temporal progenitor programme, thus delaying the generation of intermediate-born ZFHX3+ neurons, and supports the conclusion that the progenitor programme controls the temporal identity of the neurons generated.

Finally, we examined cell type specific expression (CaTS-RNA) of known temporal markers of motor neurons and oligodendrocyte precursors (OPC). In *Nfia*/*Nfib* dKO cells, we found not only a failure to upregulate markers of the OPC programme (*Olig1*, *Sox10*), but also an extended window of expression for early motor neuron markers (*Mnx1*, *Isl1*, *Lhx3*) until day 11 (Fig 6F), consistent with the chicken data (Fig. 5C). Together, these data show that perturbation of the temporal programme in progenitors changes the differentiated cell type output, consistent with the notion that the global chromatin accessibility controls cell diversity generation in the neural tube.

## Discussion

The formation of a functioning nervous system depends on the generation of the right cells in the right place, at the right time. Here we define a chronotopic mechanism for the integration of the spatial and temporal cues that control the timing and pattern of cell type specification in the vertebrate neural tube. A global temporal chromatin remodelling programme operates along both the dorsal-ventral and rostral-caudal axis of the nervous system to modify chromatin accessibility in neural progenitors. This determines the output of spatially specific gene expression programmes to govern the time and position of cell type generation. Disrupting key temporal regulators alters lineage progression and differentiated cell identities, demonstrating the temporal identity of progenitors directs cell fate decisions, as also shown in other regions of the neuraxis (Lyu et al., 2021). Together the data indicate that spatial and temporal patterning mechanisms function via distinct regulatory strategies to enable dynamic and cell type-specific gene expression over developmental time.

Previous work had identified temporal gene expression cascades directing sequential transitions in progenitor states (Sagner et al., 2021; Telley et al., 2019; Tiklová et al., 2019). Here, we demonstrate perturbing this temporal programme in progenitors affects differentiated cell identities. The early factor, Nr6a1 (GCNF), impedes the temporal progression of neural progenitors and must be downregulated for the regulatory programme to proceed. Extending the expression window of Nr6a1 alters the differentiated cell types generated in vitro and in vivo. Nr6a1 is also implicated in the temporal programme of other developing tissues, controlling the timing of Hox gene activation and rostral-caudal patterning during body axis elongation (Chang et al., 2022). Nr6a1 interacts directly with several chromatin modifiers DNMT3b, and methyl-CpG (Fuhrmann et al., 2001)(Sato et al., 2006)(Gu et al., 2011) suggesting that Nr6a1 acts as a global chromatin epigenetic regulator.

Whilst Nr6a1 promotes the early temporal phase, Sox9 and NFIA/NFIB are important for later temporal phases (Kang et al., 2012). The data show that NFIA/NFIB function via modifying chromatin accessibility and are necessary for the temporal progression of cell type specification. Loss of NFIA/NFIB affected numerous elements in progenitors that are predicted to mediate late gene induction events. The affected genes were spatially variable, which is expected if spatial determinants bind temporally available sites in a cell type-specific way. NFIA/NFIB and SOX9 regulate chromatin accessibility in several tissues (Denny et al., 2016) (Tchieu et al., 2019) (Yang et al., 2023) and recruit the chromatin remodelling complexes (Liu et al., 2001)(Vierbuchen et al., 2017; Yang et al., 2023). Whether the temporal expression of chromatin modifying and remodelling complexes is a crucial part of the temporal programme of the neural tube remains to be determined. The progressive downregulation of PRC2 in mouse cortex has been implicated in temporal progression in the cortex (Telley et al., 2019) and it is suggested to be part of an epigenetic barrier to neuronal maturation (Ciceri et al., 2024), supporting a causal role for epigenetic regulators in controlling timing.

The scale of the temporal changes in chromatin accessibility in neural progenitors was unexpected given the similarity of the chromatin landscape between spatially distinct progenitor domains (Delás et al., 2023). The dominance of differential accessibility underpinning temporal changes, compared with spatial patterning, suggests a strategy for the integration of spatial and temporal programmes. In this view, temporal factors act similarly to ‘pioneer factors’ (Barral and Zaret, 2024), altering the chromatin landscape and defining the set of available cis regulatory elements. Eviction of spatial determinants from recently closed elements and their recruitment to newly opened elements would occur in a domain-specific manner due to the spatially restricted expression of the spatial factors. This chronotopic mechanism converts shared accessibility across progenitor domains into spatial heterogeneity in transcription factor binding to determine domain and time specific gene expression. Mapping the binding patterns of temporal versus spatial factors across these elements over time and understanding how the accessibility of regulatory elements changes will further define this regulatory logic.

This strategy for integrating spatial and temporal patterning appears distinct from that operating in *Drosophila*, where STFs, expressed early in distinct progenitor cells, establish lineage-specific chromatin accessibility profiles that remain constant over time – a spatiotopic chromatin landscape (Sen et al., 2019). *Drosophila* TTFs thus generate cell type diversity by imparting a different gene expression output in each lineage, depending on the chromatin accessibility established by STF. By contrast, in the vertebrate neural tube, the opposite regulatory logic prevails: vertebrate TTFs sculpt a global temporal chromatin landscape that directs where spatially restricted TFs bind. Changes in the TTFs over time alter chromatin accessibility in all lineages, producing the chronotopic chromatin landscape. For example, TTFs such as NFIA/B establish progressive opening of regulatory elements in all progenitor domains. The differential chromatin accessibility over time results in temporally varying lineage specific gene expression and cell type generation.

What might account for this difference in regulatory logic between flies and vertebrates? In vertebrates, patterning occurs as the neural tube grows and the spatial cues, provided by morphogen gradients, act dynamically (Frith et al., 2023). Accordingly, progenitors change their expression of spatial transcription factors as the neural tube develops. This may favour a strategy in which a global temporal programme controlling chromatin landscapes across progenitors provides regulatory elements for morphogen controlled spatial cues to bind differentially over time. By contrast, *Drosophila* neuroblast lineages are established early and maintain their spatial identity throughout development (Sen et al., 2019). Perhaps this favours a strategy in which spatially distinct chromatin landscapes persist to direct differential TTF binding. Overall, the *Drosophila* and vertebrate strategies highlight alternative but effective solutions to a common problem: generating cell type diversity across space and time from multipotent progenitors.

These results have important implications for engineering specific neuron and glia subtypes using stem cells. Much attention has focused on establishing the appropriate spatial gene expression profile to reprogram cells (Chen et al., 2020; Lin et al., 2023; Nehme et al., 2018). Less regard has been given to the temporal programme. However, our data indicate that how the spatial programme is interpreted depends on the temporal chromatin state. Considering only the spatial transcriptional programme risks incomplete or misdirected reprogramming. Inappropriate temporal states may result in required regulatory elements being inaccessible, or inappropriate elements being accessible, thereby misdirecting spatial transcription factors. This might explain inefficiencies and off-target results of some current directed differentiation protocols. Our results suggest imposing the desired temporal epigenetic state followed by manipulating spatial identity might allow more rapid generation of clinically relevant cell types, such as oligodendrocyte progenitors. Overall, clarifying the regulatory logic directing progenitor temporal identity will likely improve directed differentiation and reprogramming strategies.

In summary, our study advances understanding of the regulatory logic governing neural progenitor diversity throughout the nervous system, revealing a novel chronotopic strategy for the integration of global temporal and spatial patterning. This mechanism offers a framework to conceptualise and study the molecular basis for cellular complexity of the nervous system.

## Acknowledgements

We are grateful to the Briscoe lab members for their constructive comments on the manuscript, Thomas Watson for initial protocol development and generation of Nfia/Nfib double knockout, and Chris Cheshire for establishing the TOBIAS footprinting pipeline. We thank Borzo Gharibi and Silvia Santos for generating the R1-Cas9 line, and the following Science Technology Platforms at the Francis Crick Institute for their assistance in carrying out experiments: Advanced Sequencing STP, Flow Cytometry STP, and the High throughput screening STP. This work was supported by the Francis Crick Institute which receives its core funding from Cancer Research UK (CC001051), the UK Medical Research Council (CC001051), and the Wellcome Trust (CC001051); by the European Research Council under European Union (EU) Horizon 2020 research and innovation program grant 742138 and by the Wellcome Trust (220379/D/20/Z). IZ was supported by Cancer Research UK (C157/A23459). GLMB is supported by EMBO ALTF (792-2021) and UKRI (EP/X031225/1). JCS was funded by a Boehringer Ingelheim Funds PhD Fellowship. TF is supported by a Sir Henry Wellcome Postdoctoral Fellowship (218670/Z/19/Z). AS is supported by funding from the Deutsche Forschungsgemeinschaft (German Research Foundation — Projektnummer 455354162).

## Author Contributions

M.J.D and J.B. conceived the project, interpreted data and wrote the manuscript. M.J.D. and I.Z. designed and performed experiments. G.L.M.B. performed chicken experiments. T.F. performed mouse embryo experiments. J.C.S. performed molecular cloning and perturbation experiments. M.J.D. and A.S. performed bioinformatic analyses. R.L.B. contributed to conceptualization and data interpretation. M.J. and M.H. contributed to the design of the CRISPR screen and performed experiments related to it.

## Competing interests

The authors declare no competing or financial interests.

## Materials and Methods

### Mice maintenance and husbandry

Mice of the following F1(B6xCBA), C57BL6 and outbred UKCrl:CD1 (ICR) (Charles River) mice were used for timed matings. Embryos for analyses were collected at the indicated time points following a mating, with the day of plug detection designated E0.5.

All animal procedures were carried out in accordance with the Animal (Scientific Procedures) Act 1986 under the Home Office project licence PP8527846. Animals were housed under a 12-h light, 12-h dark cycle at the Francis Crick Institute animal facility. Animals were housed in singly-ventilated cages. The relative humidity was kept at 45 to 65%. Mouse rooms and cages were kept at a temperature range of 20-24°C. Animals had 24 hour access to RO water and autoclaved pelleted food. Caging, bedding, nesting materials and enrichments were autoclaved prior to use and cages were changed routinely. A maximum of 5 adult mice were housed per cage if they were over 20 g. Animals were monitored visually daily for health concerns and once a week a full health check was carried out during a discretionary cage change.

### Chicken embryos

Fertilized chicken embryos (Gallus gallus) were obtained from Henry Stewart & Co. Ltd and were never incubated beyond two thirds of development.

### Cell lines

Wild-type experiments were performed with the mouse embryonic stem cell line HM1 (Doetschman et al., 1987) (Thermo Fisher, MES4303). The Nfia;Nfib knockout mutant mouse ES cell line (dKO) was generated into the HM1 wild-type background (Sagner et al., 2021). All cell lines were maintained at 37°C with 5% carbon dioxide (CO2).

### Generation of R1-iCas9 ES cell line

R1-iCas9 (Clone 2E9) mouse embryonic stem cell line was generated using a modified version of the Genome-CRISPTM Mouse Rosa26 Safe Harbour Gene Knock-in Kit and SH354-ROSA26 plasmid (GeneCopoeia). A Cas9 expression cassette was cloned into a FLAG tagged SH354 plasmid with a Tet-On 3G promoter system (driven by EF1a) integrated into the Rosa26 safe harbour locus in the R1 mouse embryonic stem cell line (ATCC® SCRC-1011TM Positive clones were selected using 0.5ug/ml Puromycin, followed by single cell cloning. R1-iCas9 cells were initially cultured in 2i condition (N2B27 medium supplemented with 1 µM PD0325901 (Cambridge Biosciences), 3 µM CHIR99021 (Axon) and 10 ng/ml mouse LIF (BioLegend)). Cells were adapted to grow on feeders by passaging them for five times.

### Cell culture and neural progenitor differentiation

Mouse ES cells were maintained on a ‘feeder’ layer of mitotically inactivated mouse embryonic fibroblasts (MEFs, derived and expanded in-house) in ES cell medium (Dulbecco’s Modified Eagle Medium (DMEM) Knock Out (Gibco; 10829-018) supplemented with 10% Foetal Bovine Serum (Pan Biotech; P30-2602), Penicillin/Streptomycin (Gibco; 15140122), 2 mM L-Glutamine (Gibco; 25030024), 2 mM Non-essential amino acids (Gibco, Cat No. 11140-035), and 0.1 mM 2-mercaptoethanol (Gibco; 21985-023)) with 1000 U/ml LIF (Chemicon, Int ESG1107). ES cell medium was changed every day. Cells were grown on 6-well plates or p6 dishes and passaged every second day. ES cells were washed once with PBS (Gibco; 14190-094) and then trypsinized using 0.05% Trypsin-EDTA (Gibco, 25300054) for 3 min at 37°C, spun down, resuspended in ES cell medium and 300,000 cells in 1.5 ml were seeded per well (500,000 in 4 ml per 1x p6).

One day before differentiation, CellBIND 6-well dishes (Corning) were coated with 0.1% gelatin (Sigma) overnight at 37°C. On Day 0 (D0, D – differentiation day) ES cells were washed once with PBS and dissociated using 0.05% Trypsin-EDTA (Gibco; 25300054) for 4 min at 37°C. Cells were resuspended in 10 ml ES media. For differentiation of ES cells, feeders were removed (‘panning’) by incubating ES cell + feeder suspension for 20 min at 37°C on 0.1% gelatin-coated 10 cm plates. This process was repeated once. For 11-day differentiations, 60-80,000 cells were plated onto the pre-coated CellBIND dishes (Corning) in N2B27 medium (Advanced DMEM - F12 (Gibco; 21331-020) and Neurobasal medium (Gibco; A35829-01) (1:1), supplemented with 1xN2 (Gibco; 17502001), 1xB27 (Gibco; 17504001), 2 mM L-glutamine (Gibco; 25030024), 40 μg/ml BSA (Sigma-Aldrich, Cat No. A7979-50ML), and 0.1 mM 2-mercaptoethanol) supplemented with 10 ng/ml bFGF (R&D; 100-18B). On day 2, the media was changed to N2B27 supplemented with 5 μM CHIR99021 (GSK3β inhibitor; Axon, 1386). 20-21 h later, on day 3, the media was changed to N2B27 with 100 nM RA and 10 nM of Smoothened Agonist (SAG; Calbiochem, 566660). From day 5 onwards, cells were kept in N2B27 supplemented with 10 nM SAG. From day 3 onwards, media changes were performed daily.

The differentiations described as “0 nM SAG” were performed as described above with no addition of SAG.

### Imaging-based 96-well plate CRISPR-Cas9 mutant screen

#### Preparation of custom library

96-well plates containing the crRNA Cherry-Pick Custom Library (0.1 nmol, 5 crRNA pool) from Horizon Discovery were spun down briefly and de-sealed in a Class II Laminar flow hood using aseptic technique. The library includes a pool of 5 crRNAs for each gene of interest per well. 50 µl of 1x siRNA buffer (Horizon Discovery, B-002000-UB-100) per well was added to make up a 2 µM crRNA stock. Plates were covered and rocked gently on a plate shaker for 30 min. Plates were then spun down briefly (1 min, 1000 rpm at room temperature). To prepare libraries, 10 µl of 2 µM crRNA stock was transferred to new V-bottom 96-well plates using a 96-well electronic pipette VIAFLO 96 (Integra Biosciences). 40 µl 1x HBSS buffer was added to make a 400 nM stock. 60 ml of 400 nM tracrRNA was prepared. 50 nmol tracrRNA was resuspended in 500 µl 1x crRNA to make a 100 µM stock. 240 µl of this stock was diluted in 59.76 ml of HBSS buffer. 50 µl of 400 nM tracrRNA was added per well to make 100 µl of 200 nM crRNA:tracrRNA stock. All crRNA and crRNA:tracrRNA stock 96-well plates were sealed and stored at -20°C until further use.

#### Screen differentiation

One day before transfection with CRISPR guides, Cas9 was induced in R1-iCas9 ES cells by exposure to 1 µg/ml Doxycycline (Sigma-Aldrich; D9891), which was supplemented to the ES cell medium. On the next day, flat µClear polystyrene bottom 96-well plates (Greiner, PS, F-Bottom, 655090) were coated with 70 µl matrigel (Corning, 356231, 1:50 in DMEM/F12) per well. Plates were briefly spun down for 1 min at 1000 rpm and incubated for 3 h at room temperature. ES cells were prepared as described above. During the panning steps, CRISPR guide RNA (gRNA) complex and transfection reagents were prepared.

The Edit-R CRISPR-Cas9 platform (Horizon Discovery) was used. The platform includes two components required for gene editing: A chemically synthesized trans-activating CRISPR RNA (tracrRNA), and a chemically synthesized CRISPR RNA (crRNA) designed for the gene target site of interest. For transfection, 5 crRNAs (20 µM stock) were pooled and combined with tracrRNA (20 µM stock, U-002005) to form a guide RNA (gRNA) complex (crRNA:tracrRNA) (final concentration: 10 µM). gRNA was then diluted in 1x HBSS buffer (Gibco, 14170-088) to make up 200 nM. As an example: 10 µl crRNA (2 µl of each individual crRNA), 10 µl tracrRNA were combined (20 µl of gRNA). 20 µl of gRNA was diluted into 1 ml HBSS. In parallel, negative, and positive controls were prepared. Non-targeting control (Control #1, 5 nmol stock, Horizon Discovery, U-007501-01-05) was used as negative control. Pool of 5 crRNAs against Nfia was used as positive control.

Matrigel was removed and plates were washed once with 200 µl PBS per well. 10 µl of the gRNA mix was added per well on the coated 96-well plates. 10 µl of diluted transfection reagent (DharmaFECT4, Horizon Discovery, T-2004-01, ∼0.15-0.2 µl per 10 µl Opti-MEM I Reduced Serum Medium (Thermo Fisher; 31985062)) was added.

R1-iCas9 cells were counted and seeded in 80 µl N2B27 medium with 10 ng/ml bFGF (R&D; 100-18B) and 1 µg/ml Doxycycline in each well. The final volume per well is 100 µl with a gRNA final concentration of 20 nM. For most experiments, 4500 cells/well were added (for the non-transfected and non-targeting controls 2500 cells/well). Per plate, at least 3 technical replicates for each targeted gene and both controls were included. Plates were rocked side-to-side to distribute cells evenly and left to sit for 5 min. Plates were moved to incubators at 37°C. Gene-edited cells were differentiated according to the previously described *in vitro* protocol.

#### Staining and readout

On day 11, plates were washed once with PBS (100 µl/well) and fixed with 4% paraformaldehyde for 15 min at room temperature. PFA was removed and plates were washed once with 200 µl PBS. 60 µl primary antibodies (anti-rabbit NFIA, 1:1000; anti-goat SOX2, 1:500) in blocking solution (PBS, 0.1% BSA and 0.1% Triton X-100) were added to each well and incubated overnight at 4°C. The next day, cells were washed once for 15 min with PBST (PBS supplemented with 0.1% Triton X-100), and secondary antibodies (Alexa 488 anti-rabbit, 1:1000; Alexa 647 anti-goat, 1:1000 in blocking solution) were added. Plates were incubated for 2-3 h at room temperature protected from light. Plates were washed once with PBS and DAPI (1:1000) for 5 min, followed by a second wash for 5 min. 200 µl PBS was added to each well, plates were sealed and stored at 4°C until imaging. Imaging was carried out within a week of staining.

#### Image acquisition and processing

Plates were imaged on the CellInsight CX7 Pro HCS Platform (Thermo Fisher) using the HCS Studio V6.6.2 software and analysed using the Cellomics Compartmental Analysis V4 (Thermo Fisher). Images were acquired at 20x (Olympus 20X/0.7NA) in ‘widefield mode’ and scanned for 25 fields of view per well using laser-based autofocus. The following channels were used: Channel 1, 647 nm (Red) SOX2; Channel 2, 488 nm (Green) NFIA; Channel 3, 647 nm (SOX2 long exposure). SOX2 positive objects were detected and segmented in Channel 1 and used as an estimator of cell number (‘ValidObjectCount’). Channel1 masks were applied to Channel 2 and 3 acquired images and the average intensity per object measured for each channel and summarised as a mean per well (‘MEAN_CircAvgIntenCh2’). The full CRISPR-Cas9 screen was performed twice (biological replicates). For each screen, three technical replicates per gene of interest were used. The processing of raw data was performed using the in-house pipeline based on the celHTS2 pipeline. In brief, non-targeting control (NT) quantifications were used for normalisation across all samples. The average values (intensity, valid object count etc.) of all NT samples were set to 100. The mean normalised intensity of all three technical replicates per gene mutated was calculated. Full downstream analysis can be found at https://github.com/MJDelas/temporal-neural.

#### Differentiation timecourse analysis by flow cytometry

Selected hits from the 96-well CRISPR screen were validated in a 12-well secondary screen using flow cytometry. Differentiations were performed in matrigel-coated CellBIND 12-well plates (Corning). CRISPR reaction volumes were calculated relative to the 96-well plate CRISPR mutant screen. 45,000 cells per well were seeded in 500 µl total volume (with a crRNA/tracrRNA complex final concentration of 20 nM). Samples were collected at day 7, 9 and 11 of differentiation. Cells were washed twice with PBS, accutased and transferred from 12-well plates into V-bottom 96-well plates (Corning, 3897). Cells were spun down at 1000 rpm for 4 min and resuspended in PBS plus LIVE/DEAD^TM^ Fixable Dead Cell Stain Near-IR fluorescent reactive dye (Thermo Fisher; L34976) and incubated for 30 min protected from light. Cells were then fixed with 4% PFA for 15 min, washed once with PBS/0.5%BSA and processed for Flow cytometry staining.

### Flow cytometry of intracellular markers

#### Sample collection

Cells were washed three times with PBS before dissociation with Accutase (Gibco, A1110501) for 5 min at 37°C. Cells were collected in ES cell medium, spun down at 400xg for 4 min and resuspended in PBS. Cell suspensions were supplemented with LIVE/DEAD^TM^ Fixable Dead Cell Stain Near-IR fluorescent reactive dye (Thermo Fisher, L34976), according to the manufacturer’s instructions (1 µl dye/1 ml PBS) and incubated for 30 min on ice protected from light. Cells were spun down at 1,000xg for 4 min and fixed with 4% PFA (Thermo Fisher, Cat. No. 28908) for 15 min on ice. After washing once with PBS, cells were resuspended in PBS + 0.5% BSA and stored at 4°C.

#### Staining

Between 1 and 2×10^6^ cells were used for flow cytometry analysis. Cells was transferred into 1.5 ml DNA LoBind® Tubes (Eppendorf) and incubated with 100 µl primary antibodies in blocking solution (PBS + 0.1% Triton X-100 + 1% BSA) overnight at 4°C or if conjugated antibodies were used, samples were incubated for 1-2 hr at RT. The next day, cells were spun down at 4000 rpm for 4 min and washed once with PBS before adding secondary antibodies together with conjugated antibodies for 1-2 hr at room temperature. Cells were washed once with PBST, spun down and resuspended in 300 or 500 µl PBS with 0.5% BSA. Flow cytometry was performed using a Becton Dickinson LSRFortessa™ Cell Analyzer (BD Biosciences) recording 10,000 SOX2-positive cells as stopping gate. Analysis of data was performed using the FlowJo software (Version 10.8.2, Becton Dickinson).

### Immunohistochemistry of embryo sections

Mouse or chicken embryos were fixed in 4% paraformaldehyde for 2 h at room temperature, cryoprotected by equilibration in 0.12 M Sodium Phosphate buffer containing 15%w/v sucrose solution overnight at 4°C. Tissues were subsequently mounted in Sodium Phosphate buffer containing 15% w/v sucrose and 7.5% w/v Gelatin and were flash frozen in Isopentane. Cryosectioning (14 μm sections) was performed on a Cryostat LEICA CM3050S using SuperFrost slides (Thermo Fisher, 10149870).

Immunostaining was performed as previously described (Delás et al., 2023), using the following antibodies. Mouse: mouse anti-NKX6.1 (1:100, DSHB, F55A10), rabbit anti-NFIA (1:1000, Atlas Antibodies, HPA008884), goat anti-OLIG2 (1:500, R&D Systems, AF2418). Chicken: rabbit anti-Olig2 (1:500, EMD Millipore, AB9610), sheep anti-Zfhx3 (1:400, R&D, AF7384), goat-Isl1 (1:500, R&D, AF1837), mouse-Sox2 (1:500, Santa Cruz, sc-365823), rat anti-RFP (1:500, Chromotek, 5f8). Secondary antibodies used were from ThermoFisher conjugated with AlexaFluor and were all used at 1:500 concentration, except AlexaFluor 647 conjugated antibodies that were used 1:1000. Sections were imaged with a Leica SP8 confocal microscope and images processed with Fiji.

### Plasmid generation

To generate the Nr6a1 chicken expression plasmids, we cloned using Golden Gate an IDT gblock containing the ORF of Nr6a1 (Gencode Transcript: ENSMUST00000076275.10), where PaqCI cut sites were removed by synonymous mutation, into an ampicilin resistant backbone, downstream of an Ef1a promoter and upstream of a T2A-mScarlet3 fusion (Addgene 189753), and an SV40 terminator. Our control vector is identical to this but lacking the Nr6a1-T2A, cloned in the same manner. The constructs were transformed using MAX Efficiency™ Stbl2™ Competent Cells (Thermo Fisher 10268019) and were grown in 100mL LB shaking for 16h at 37°C, before maxi-prepping using the QIAGEN Plasmid Plus Maxi Kit (12963).

### Plasmid backbone and virus generation

To generate Nr6a1 lentivirus expression vectors, we amplified by PCR the Ef1a to SV40 inclusive segments of the chicken overexpression plasmids to include Esp3I Golden Gate adaptors using Phusion Flash (Thermo Fisher; F548L), into a lentivirus backbone derived from (Delás et al., 2017). Constructs were transformed using MAX Efficiency™ Stbl2™ Competent Cells (Thermo Fisher10268019) and maxi-prepped as above. To generate lentivirus, HEK293T were plated at 1.5×10^6^ per 6cm plate. The next day, the media was changed (3.5mL). A mixture of 3rd generation packaging plasmids (0.625μg of CMV-Rev, 1.25 μg pMDLg and 0.9 μg VSV-G per 5 μl) and 2.28 μg of the transfer plasmid was vortexed with 11.9uL X-tremeGENE™ HP DNA Transfection Reagent (Roche; 6366244001) and 360 μl OptiMEM (GIBCO; 31985062). This was added dropwise on top of the cells. The media was changed 16h later to N2B27 and collected 30hrs later. The media was filtered through 0.45 micron filter and frozen at -80°C in aliquots.

### RNA extraction and RT-qPCR

Cells were washed twice with PBS and 350 µl RLT lysis buffer (Qiagen, 1015762) was added directly to the dish (e.g. p3.5/35 mm or 1 well of 6-well plate). After 5 min lysed cells were collected and transferred to a 2 ml RNase-free Eppendorf tube and stored at -20°C for no longer than 2 weeks until further processing. Total RNA was extracted using RNeasy Mini Kit with DNAse digest (Qiagen, Cat. No. 74106) according to the manufacturer’s instructions and stored at -80°C. 1.5 μg of RNA was reverse transcribed into cDNA using SuperScript III first strand synthesis system (Invitrogen 18080-051) using random hexamers. qRT-PCR was performed in 384-well plates (reaction volume 10 µl) using Platinum™ SYBR™ Green qPCR SuperMix-UDG (Invitrogen A25742) on either the QuantStudio 5 or 12K Flex Real-Time PCR system (ThermoFisher Scientific). Expression values (CT values) were normalised against β-Actin. All experiments were performed in technical duplicates, biological duplicates or triplicates for each time point analysed. Primers were designed using NCBI tool Primer BLAST unless stated otherwise.

### CaTS-ATAC-seq (Crosslinked and TF-Sorted ATAC-seq)

CaTS-ATAC was performed as previously described (Delás et al., 2023) with in-house produced Tn5. A dead cell removal step was introduced to enrich for live cells prior to fixation and transposition. The details are as follows: at the differentiation timepoint of collection, cells were washed once with PBS and dissociated with Accutase (Gibco, A1110501) for 5 min at 37°C. Cells were collected in 1.5 ml LoBind Eppendorf tubes (Eppendorf cat. #Z666548) and spun down at 400xg for 4 min. Cell suspensions were supplemented with LIVE/DEAD^TM^ Fixable Dead Cell Stain Near-IR fluorescent reactive dye (Thermo Fisher, L34976), according to the manufacturer’s instructions (1 µl dye/1 ml PBS for 1×10^6^ cells) and incubated for 30 min on ice protected from light. During the incubation, the autoMACS^®^ Pro Separator (Miltenyi Biotec) was set-up according to the manufacturer’s instructions. After the 30 min cell suspensions were divided into 15 ml falcon tubes and spun down for 10 min at 400xg at 4°C. Cells were resuspended, pooled into one tube, and counted. Cells were magnetically labelled with Dead Cell Removal Microbeads from the Dead Cell Removal Kit (Miltenyi Biotec, 130-090-101). For 1.0×10^7^ cells, 100 µl beads were used. Cells and microbeads were mixed and incubated for 15 min at room temperature. If necessary, 1x Binding Buffer provided by the kit was added to the cell suspension to reach a minimum volume of 500 μl for separation. The samples were run through the autoMACS® Pro Separator (Miltenyi Biotec) using the ‘Depl05’ programme. The microbeads label dead cells, which are retained in the separation columns. Unlabelled live cells run through the column and were collected in a 15 ml falcon tube in PBS. Live cells were count and spun down for 4 min at 400xg at 4°C. Cell pellet was resuspended in 300 µl of PBS in 1.5 ml LoBind Eppendorf tubes. Cells were then fixed, transposed, stained for viability and intracellular markers (anti-SOX2-V450, anti-Nkx6.1-PE, goat anti-Olig2, rabbit anti-Nfia) and sorted. DNA of samples was isolated using the DNA Clean & Concentrator-5 kit (Zymo; D4013) as per manufacturer’s instructions.

### CaTS-RNA-seq (Crosslinked and TF-Sorted RNA-seq)

Samples for CaTS-RNA-seq were collected in the same way as for CaTS-ATAC-seq until the fixation step. From there on all buffers were supplemented with 1:100 RNasin Plus RNase inhibitor (Promega, Cat. No. N2615).

#### Glyoxal fixation

The fixation protocol was adapted from published work (Channathodiyil and Houseley, 2021). To prepare ∼4 ml of 3% glyoxal solution, 2.835 ml of RNase-free water was combined with 0.789 ml 100% (v/v) EtOH, 0.03 ml acetic acid and 0.313 ml 40% Glyoxal (Sigma-Aldrich, Cat. No. 50649-100ML). All steps were performed on ice and reagents were cooled before mixing. The solution was adjusted to pH 4-5 (with 1M NaOH) if necessary. The solution was always kept on ice and stored for up to 3 days at 4°C. All samples were fixed and stained on the same day. To minimise RNase activity, cells were kept on ice and only pre-chilled reagents were used.

Cells were resuspended in 50 µl PBS + 0.5% BSA first, followed by adding 300 µl 3% glyoxal. Cells were fixed for 15 min with slow side-to-side rocking on ice protected from light. 25 µl of 2M glycine was added for quenching and incubated for 5 min. 1 ml PBS + 0.5 % BSA was added to stop the fixation, and cells were spun down at 2000xg for 5 min at 4°C. 1-2×10^6^ cells were prepared for Flow cytometry staining and resuspended in 100 µl PBS + 0.5% BSA + 0.3% Triton X-100. Cells were incubated on ice for 30 min. Staining was performed as previously described and processed in the same way as described for CaTS-ATAC-seq. 10-15,000 cells of each desired population were sorted into 300 µl TRIzol reagent (Invitrogen, Cat. No. 15596026) and stored at -80°C until further processing.

#### TRIzol RNA extraction and library preparation

All steps were performed at room temperature unless otherwise stated. For TRIzol extraction, Phasemaker tubes (Thermo Fisher A33248) were used according to the manufacturer’s instructions. 300 µl of sorted cells in TRIzol were mixed and transferred to Phasemaker tubes and incubated for 5 min at room temperature. 200 µl chloroform per 1 ml TRIzol used for lysis was added and mixed vigorously. Samples were incubated for 10 min and spun for 5 min at 12,000×g and 4°C. The upper phase was transferred to a new RNase-free tube. 8 µg RNase-free GlycoBlue (stock 15 mg/ml) was added to each sample and briefly mixed to see the pellet easier. 150 µl of 100% isopropanol was added (500 µl per initial 1 ml TRIzol). The solution was incubated for 10 min, then centrifuged for 10 min at 12,000xg at 4°C and supernatant was carefully discarded. The pellet was washed in 1 ml of 75% EtOH per 1 ml TRIzol reagent used for lysis. The sample was then centrifuged for 5 min at 7500xg at 4°C. The pellet was air dried for 5-10 min to remove residual EtOH. To solubilise the pellet, it was resuspended in ∼17 µl of RNase-free water. The resuspended RNA was incubated in a heat block for 10 min at 55°C. RNA was stored at -80°C until further processing.

After analysis RNA integrity using the Agilent Bioanalyser, paired-end library preparation was performed using the SMART-Seq HT kit (Takara, Cat. No. 634437) followed by Nextera XT DNA Library Preparation Kit (Illumina, Cat. No. FC-131-1096). Indexed libraries were pooled and sequenced on an Illumina HiSeq 4000 flow cell configured to generate 100 bp of pair-ended (PE) data (Number of reads per sample: 25 million pairs).

### Bulk RNA-seq from CRISPR-transfected mutants

Samples were collected at day 7, 9 and 11 of differentiation and transferred from 12-well plates into 96-well plates (see above). Cells were spun down and pellets stored at -80°C. Total RNA was extracted using RNeasy Mini Kit with DNAse digest (Qiagen, Cat. No. 74106) according to the manufacturer’s instructions and stored at -80°C. Libraries were prepared using the NEBNext Ultra II Directional kit(NEB, Cat no E7760L) with ribodepletion (Qiagen QIAseq FastSelect -rRNA HMR Kit, Cat. No. 334386), following manufacturer’s instructions.

### Bulk ATAC-seq from lentiviral overexpressing sorted cells

Differentiations were transduced at day 5 with lentivirus overexpressing either *Ef1a:mScarlet3* or *Ef1a:Nr6a1-T2A-mScarlet3*. Three independent differentiation wells were transduced as biological replicates. At day 9, live mScarlet3+ positive cells were sorted and 50,000 cells per replicate were used for ATAC-seq library preparation. Bulk ATAC-seq on live cells was performed as previously described (Delás et al., 2023).

### Chicken embryos and electroporation

Fertilized chicken embryos (*Gallus gallus*) were obtained from Henry Stewart & Co. Ltd and kept in humidified incubators at 38°C up to the desired Hamburger and Hamilton (HH) stage. Eggs were incubated for 40-48 h to reach HH11-HH12 stages and injected in the developing neural tube via a pulled glass capillary (Harvard Apparatus, EC1 64-0766) with plasmid solutions mixed with SYBR Green (Thermo Fisher, S7563). *Ef1a:mScarlet3* or *Ef1a:Nr6a1-T2A-mScarlet3* plasmids were injected at a final concentration of 150-300 ng/µl, according to the concentration that ensured consistent mScarlet3 expression. Electroporations were performed as previously described (Balaskas et al., 2012) by passing three 24 V pulses for 50 ms each every 150 ms with an Electro Square Porator ECM830. The embryos were then incubated until stage HH23 (day 4) or HH25 (day 5) and screened for transfection efficiency (i.e., presence of mScarlet3 fluorescence within the neural tube). Healthy and correctly transfected embryos were dissected, by selecting brachial regions of the trunk, and processed for immunohistochemistry.

### ATAC-seq processing

Data was processed using the nf-core atacseq pipeline (https://nf-co.re/atacseq) with the following options: --genome mm10 --skip_diff_analysis --min_reps_consensus 2 -r 1.2.1.

### RNA-seq processing

Data was processed using the nf-core rnaseq pipeline with the following options: --genome mm10 --aligner star_salmon --skip_biotype_qc --deseq2_vst -r 3.5.

### ATAC-seq differential expression analysis

Read counts within the consensus intervals generated by featureCounts were used as input for DESeq2 (Love et al., 2014). Principal Component Analysis was performed using the top 30000 most variable elements and coloured by different sample metadata.

To assess the cell type specific accessibility for the same NP cell types at different timepoints, or the differential accessibility between timepoints for the same cell type, pairwise differential expression was performed. The number of differentially accessible intervals plotted (fold change > 2, basemean > 100, p-adj < 0.01) as a bubble plot.

To assemble the subset of highly-confident global temporally dynamic elements, differentially accessible (thresholds as above) elements in all three cell types (but in any number of temporal comparisons) were selected.

Elements affected by Nfia/Nfib double knockout were identified using the same thresholds in KO vs wildtype comparisons.

All significantly differentially accessible elements (no fold change cut off) between control and Nr6a1 overexpression are plotted coloured by p-adjusted value.

### ATAC element clusters by kmeans

Variance stabilized transformed data generated using DESeq2 were used as input to identify clusters of elements with the same dynamics. Clustering was performed using kmeans with a high number of centres, 30, and subsequently re-grouping clusters of very similar dynamics using hclust and target of 7 final clusters. This was chosen as independent iterations resulted in reproducible clusters and dynamics.

### RNA-seq differential expression analysis

The read counts from salmon were used as input for DESeq2. The threshholds used to consider a gene differentially expressed were filter(padj < 0.05 & abs(log2FoldChange) > 1 & baseMean > 80). To assemble the subset of global dynamic genes, we selected those differentially expressed in all cell types but in any number of temporal comparisons. To assemble the list of genes temporally dynamic but also cell type specific, we required genes to be differentially expressed either (i) at more than one timepoint and in at least one cell type comparison or (ii) in at least one timepoint and in two cell type comparisons.

The effects of the different mutations were examined by performing differential expression between control and KO at each timepoint.

### GO enrichment analysis of temporal clusters

Global temporally dynamic genes were clustered using ComplexHeatmap (Gu et al., 2016) into 6 reproducible clusters. GO enrichment analysis was performed using clusterProfileR (Yu et al., 2012).

### Association of genes with candidate elements

The genes identified as temporally dynamic and also differentially expressed between cell types were used as input for figR’s function runGenePeakcorr with windowPadSize = 500000 (Kartha et al., 2022). The gene to peak correlation assignments were performed independently for each cell type. For each element-gene pair, the most significant, or most correlated in case of equal pval, cell type values were kept. The accessibility of elements assigned to input genes with pvalz < 0.1, rObs > 0 was plotted. For visualization purposes, genes were clustered and their assigned genes plotted in the same order.

### Reanalysis of scATACseq across the neuraxis

In vivo spinal cord single cell data (Shu et al., 2022) was downloaded as processed bigwig files and the coverage for the spinal cord temporal elements was plotted using deeptools plotHeatmap (Ramírez et al., 2016) function. Data from other regions (Lyu et al., 2021; Bella et al., 2021; Sun et al., 2024; Sarropoulos et al., 2021) were downloaded as mapped files and ArchR (Granja et al., 2021) was used to filter the cells from the appropriate clusters and export normalize bigwig files. Deeptools plotHeatmap or plotProfile were used for data visualization.

### Data and availability

All the sequencing data asscociated with this study is available at GEO SuperSeries GSE264172. The code is available at: https://github.com/MJDelas/temporal-neural

## Supplemental figures

**Supplemental Figure 1.**
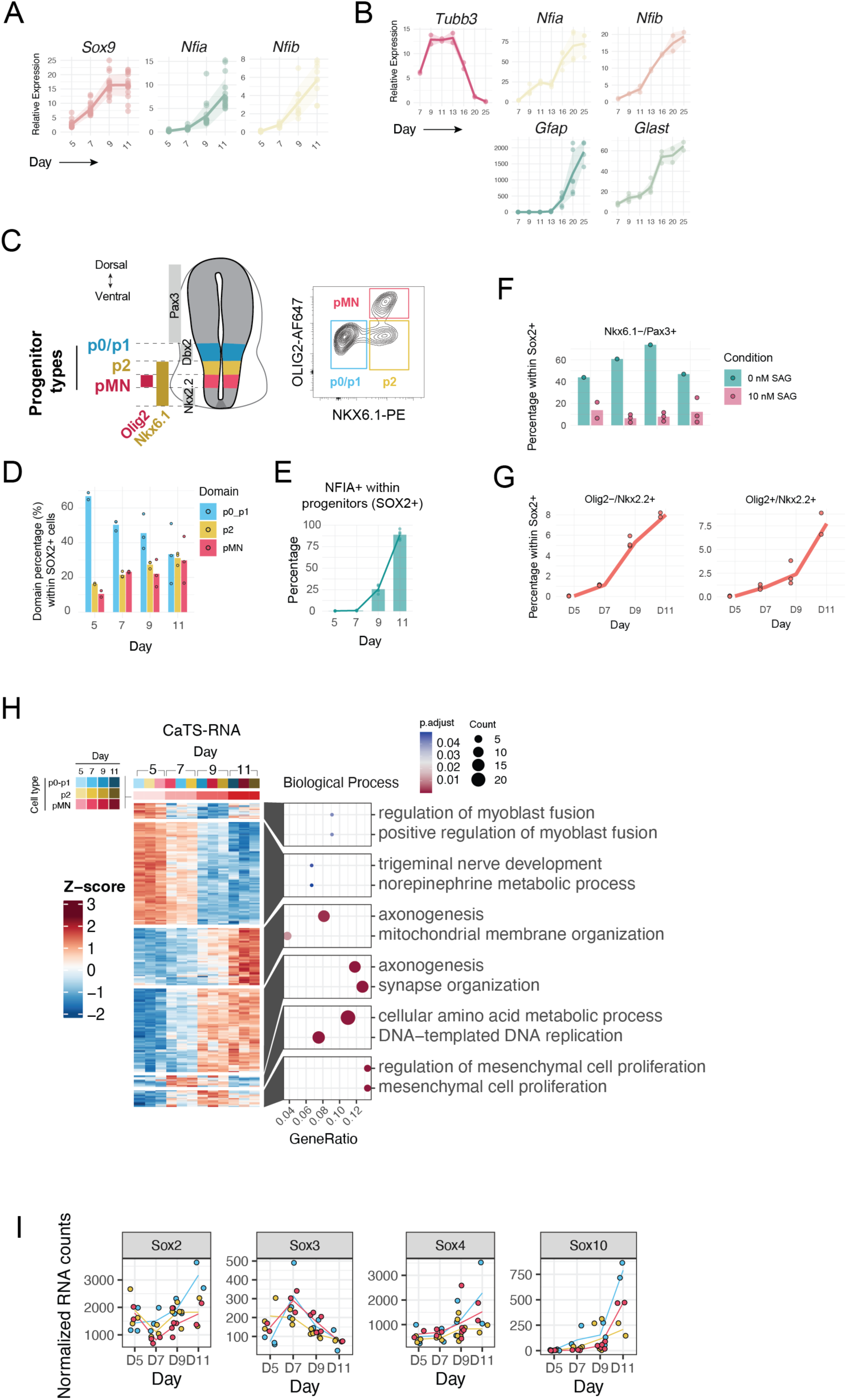
(A) RT-qPCR highlighting key genes in the temporal progression to day 11, and (B) to day 25. (C) Diagram of the spatial domains analysed in this study and the markers for the cell types of interest and the neighbouring domains (lef), and example flow cytometry plot for the cell types of interest (right). (D) Proportions of cell types (spatial domains) obtained in the cellular model over the time course progression, as measured by flow cytometry. (E) Proportion of NFIA in the cellular model as measure by flow cytometry. (F) Proportion of cells dorsal to p0 (PAX3+) was very low in the conditions used for this study (10 nM SAG). Proportions obtained with 0 nM SAG are shown for comparison. (G) Proportion of cells ventral to pMN (NKX2.2+) were only detected at very low levels at the later timepoints (left). NKX2.2 is known to be induced as part of oligodendrocyte progenitor specification and some co-expression of Olig2 and Nkx2.2 is present as expected. (H) Global gene expression over time shows differential gene ontology enrichment for early clusters (e.g. nerve development) versus later cluster (e.g. axonogenesis). (I) Normalised RNA expression of additional genes that could contribute to the SOX/1 footprint.

**Supplemental Figure 2.**
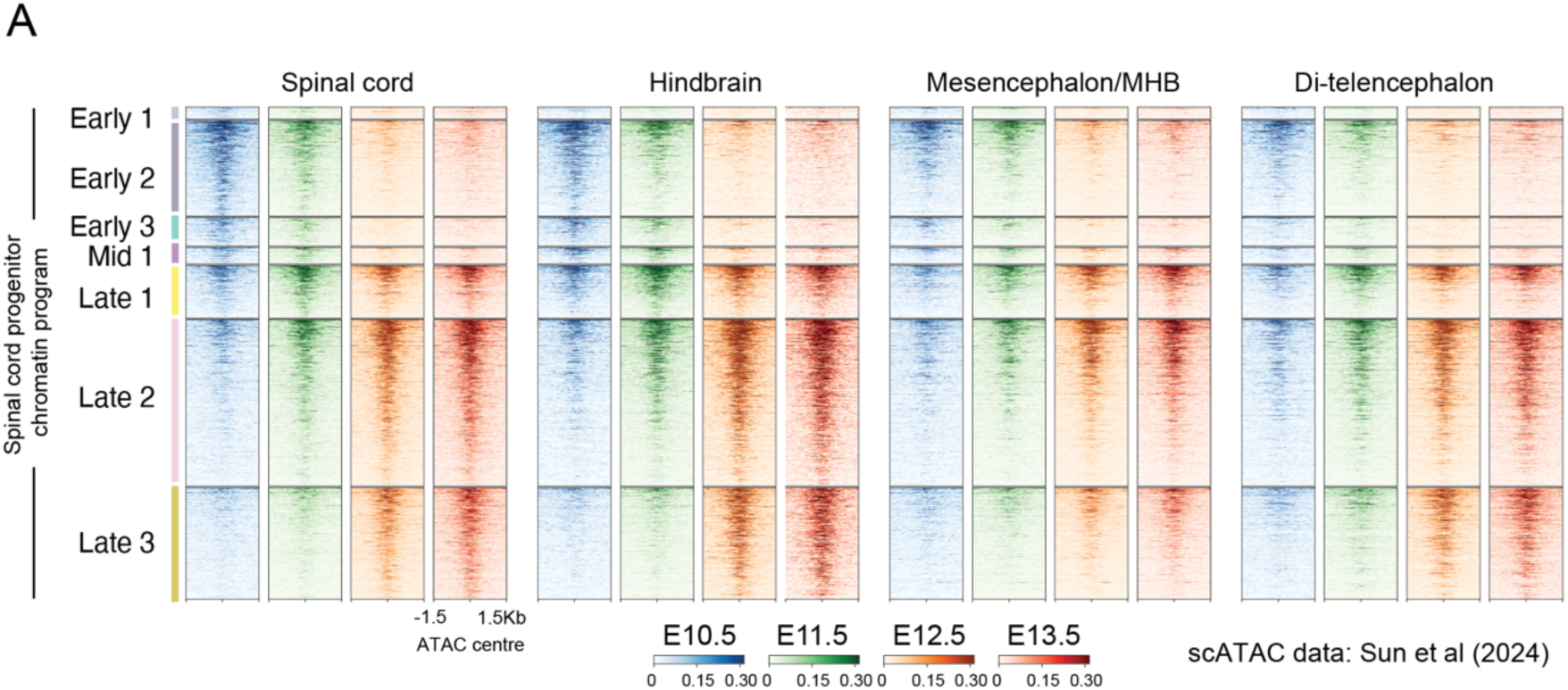
(A) Heatmap coverage plots of accessibility at the temporal chromatin elements for cells from the spinal cord, hindbrain, mesencephalon and di-telencephalon. The elements ploted are those from the spinal cord progenitor temporal programme (Fig 2A; Shu et al., 2022). Data are from (Sun et al., 2024)

**Supplemental Figure 3.**
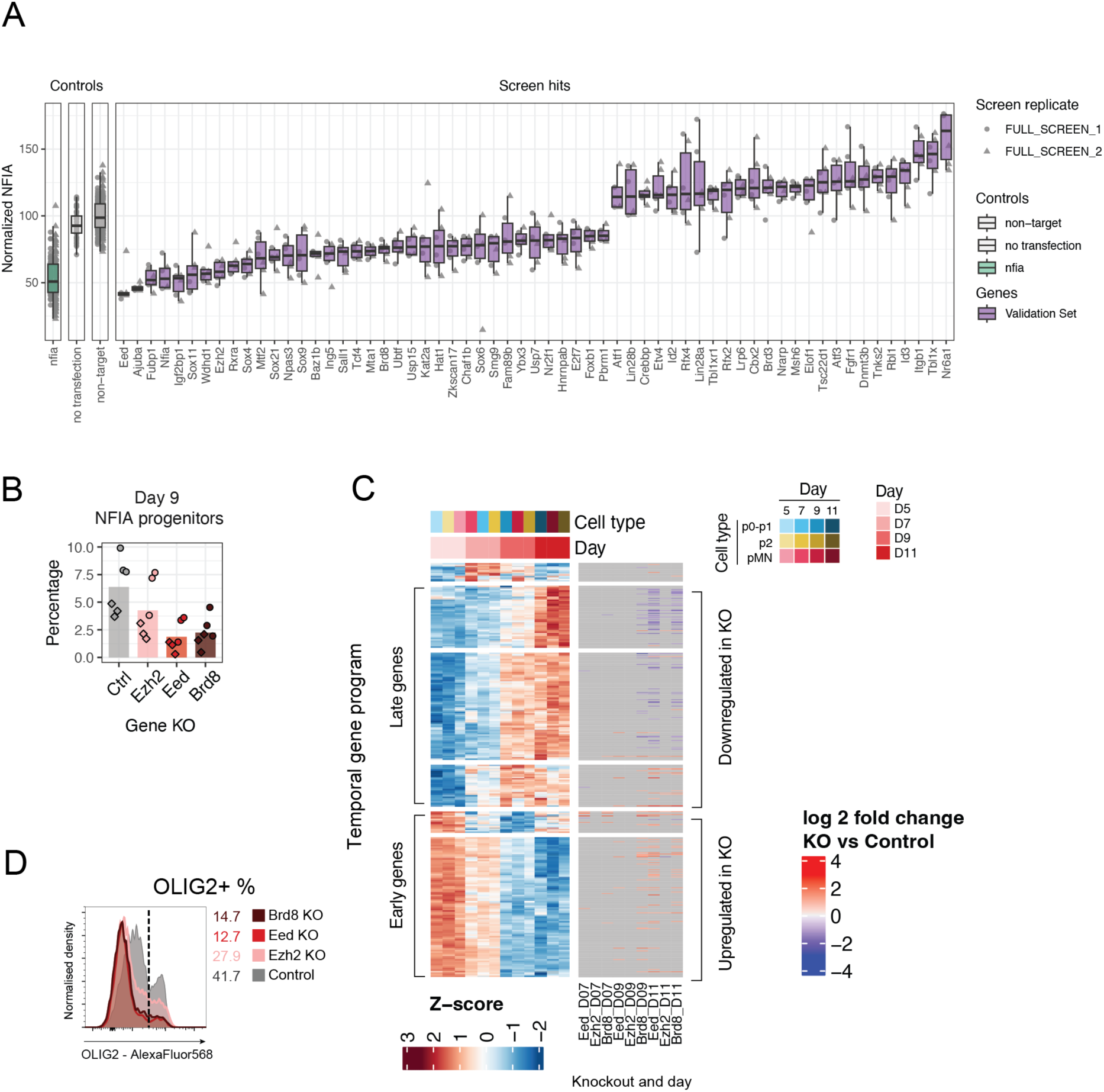
(B) Normalised NFIA intensity for all technical replicates in each of the two biological replicates of the CRISPR screen for all controls and genes selected as hits. (C) Flow cytometry staining showing the proportion of NFIA + cells within the SOX2 NPs for each of the indicated knockouts at day 9. (D) Knockout of either Eed, Ezh2 or Brd8 results in changes in genes identified as part of the temporal program, with late genes being downregulated and early genes being upregulated in knockout compared to control. (E) The proportion of OLIG2+ progenitor cells is reduced when *Ezh2, Eed* or *Brd8* are knocked out compared to non-targetting controls. Data shown for day 11.

**Supplemental Figure 4.**
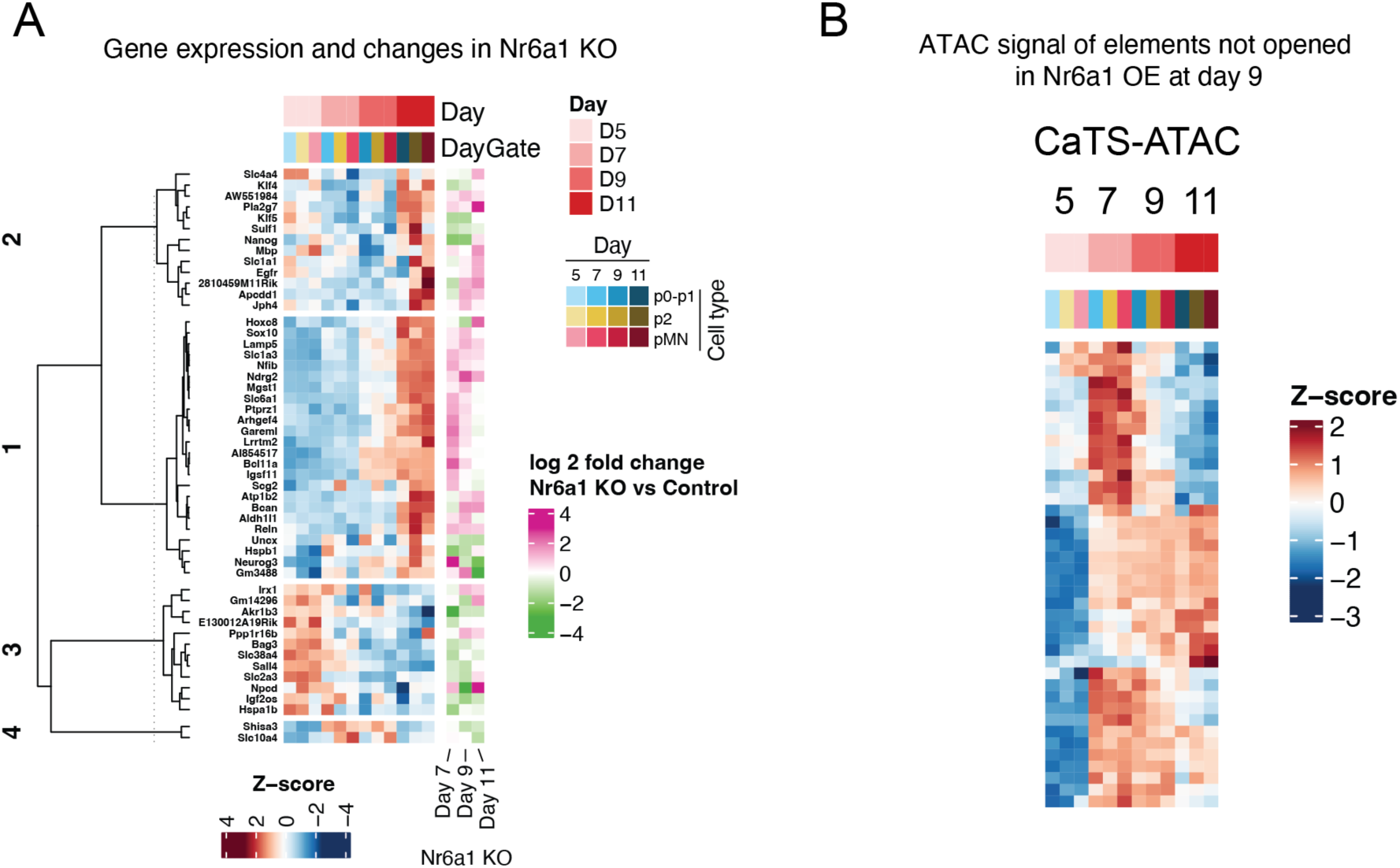
(A) Heatmap of genes affected by Nr6a1 KO showing the expression during normal differentiation (left) and the changes observed in Nr6a1 KO compared to control (right). (B) Accessibility z-scores of the elements during the normal differentiation identified as only opened in control but not in Nr6a1 overexpression.

**Supplemental Figure 5.**
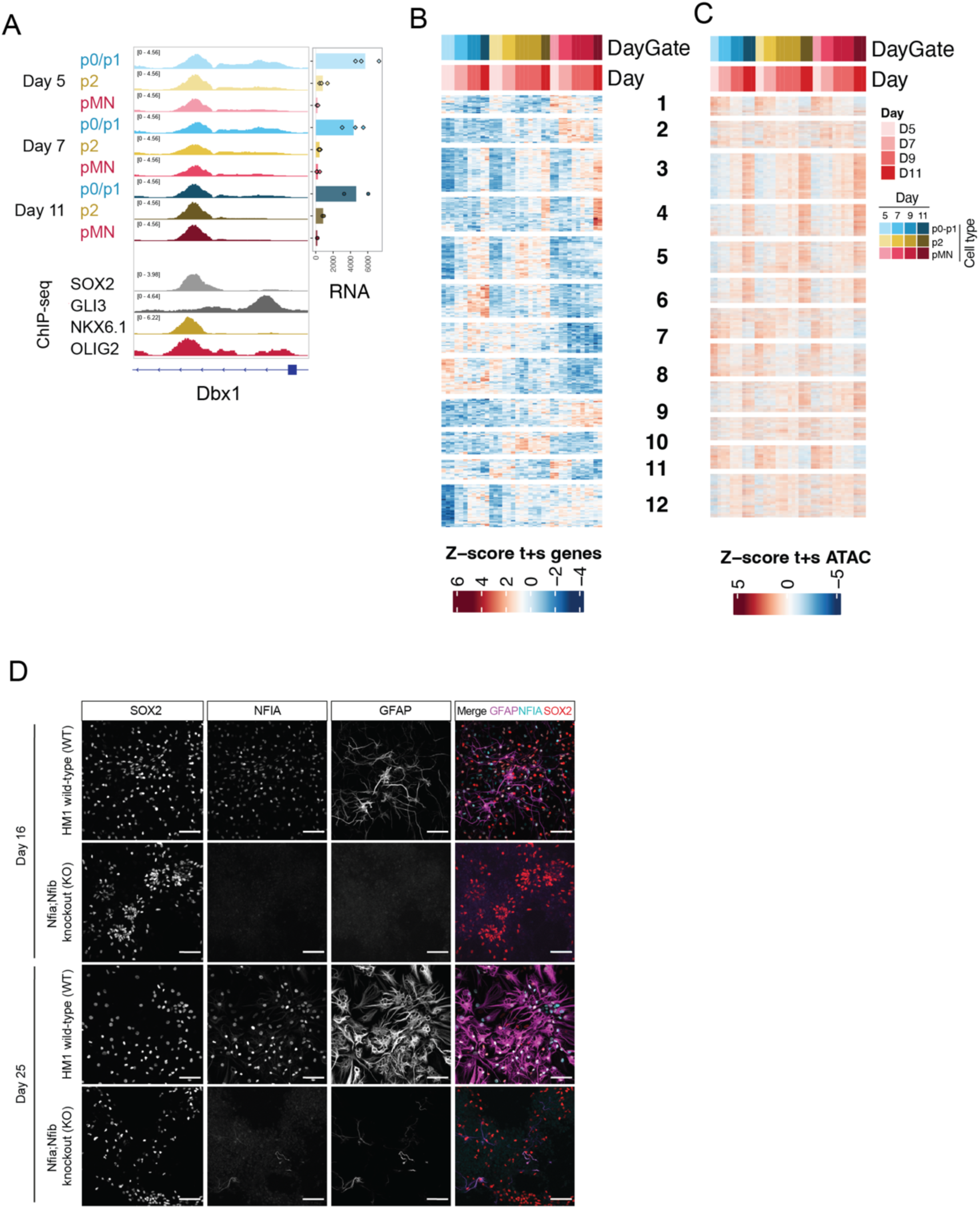
A) ATAC-seq accessibility and RNA expression of Dbx1 in the different cell types and timepoints shows global accessibility across cell types despite the cell type specific gene expression. ChIP-seq shows the element is bound by cell type specific repressors, global activators and effects of the Shh pathway. (B) Genes identified as being dynamically regulated over time and differentially expressed between cell types, clustered into 10 groups of different expression dynamics. (C) Predicted regulatory elements associated with the genes in (B) show dynamic temporal accessibility but always consistent across all cell types. (D) The ability of Nfia/Nfib dKO to upregulate the astrocytic marker GFAP is greatly impared during in vitro differentiation. Scale bar 60 µm.

**Table S1**

Oligonucleotides used in this study

**Table S2**

Antibodies used in this study

**Table S3**

Temporal genes across all domains (Fig 3, Fig S1H)

**Table S4**

Cell type specific and temporally dynamic genes (Fig 5B, S4B)

## Notes

https://www.ncbi.nlm.nih.gov/geo/query/acc.cgi?acc=GSE264172

### Competing Interest Statement

The authors have declared no competing interest.

### Summary of Updates

New figure 2 added demonstrating chromatin temporal programme is conserved across the central nervous system.

## References

Amberg, N., Pauler, F.M., Streicher, C., Hippenmeyer, S., 2022. Tissue-wide genetic and cellular landscape shapes the execution of sequential PRC2 functions in neural stem cell lineage progression. Sci. Adv. 8, eabq1263. 10.1126/sciadv.abq1263

Balaskas, N., Ribeiro, A., Panovska, J., Dessaud, E., Sasai, N., Page, K.M., Briscoe, J., Ribes, V., 2012. Gene Regulatory Logic for Reading the Sonic Hedgehog Signaling Gradient in the Vertebrate Neural Tube. Cell 148, 273–284. 10.1016/j.cell.2011.10.047

Barral, A., Zaret, K.S., 2024. Pioneer factors: roles and their regulation in development. Trends Genet. 40, 134–148. 10.1016/j.tig.2023.10.007

Bayraktar, O.A., Doe, C.Q., 2013. Combinatorial temporal patterning in progenitors expands neural diversity. Nature 498, 449–455. 10.1038/nature12266

Bella, D.J.D., Habibi, E., Stickels, R.R., Scalia, G., Brown, J., Yadollahpour, P., Yang, S.M., Abbate, C., Biancalani, T., Macosko, E.Z., Chen, F., Regev, A., Arlotta, P., 2021. Molecular logic of cellular diversification in the mouse cerebral cortex. Nature 595, 554–559. 10.1038/s41586-021-03670-5

Bentsen, M., Goymann, P., Schultheis, H., Klee, K., Petrova, A., Wiegandt, R., Fust, A., Preussner, J., Kuenne, C., Braun, T., Kim, J., Looso, M., 2020. ATAC-seq footprinting unravels kinetics of transcription factor binding during zygotic genome activation. Nature Communications 11, 4267–11. 10.1038/s41467-020-18035-1

Briscoe, J., Pierani, A., Jessell, T.M., Ericson, J., 2000. A homeodomain protein code specifies progenitor cell identity and neuronal fate in the ventral neural tube. Cell 101, 435–445.

Butt, S.J.B., Fuccillo, M., Nery, S., Noctor, S., Kriegstein, A., Corbin, J.G., Fishell, G., 2005. The Temporal and Spatial Origins of Cortical Interneurons Predict Their Physiological Subtype. Neuron 48, 591–604. 10.1016/j.neuron.2005.09.034

Cadwell, C.R., Bhaduri, A., Mostajo-Radji, M.A., Keefe, M.G., Nowakowski, T.J., 2019. Development and Arealization of the Cerebral Cortex. Neuron 103, 980–1004. 10.1016/j.neuron.2019.07.009

Cepko, C., 2014. Intrinsically different retinal progenitor cells produce specific types of progeny. Nature Publishing Group 15, 615–627. 10.1038/nrn3767

Chang, Y.-C., Manent, J., Schroeder, J., Wong, S.F.L., Hauswirth, G.M., Shylo, N.A., Moore, E.L., Achilleos, A., Garside, V., Polo, J.M., Trainor, P., McGlinn, E., 2022. Nr6a1 controls Hox expression dynamics and is a master regulator of vertebrate trunk development. Nat. Commun. 13, 7766. 10.1038/s41467-022-35303-4

Channathodiyil, P., Houseley, J., 2021. Glyoxal fixation facilitates transcriptome analysis after antigen staining and cell sorting by flow cytometry. PLoS ONE 16, e0240769. 10.1371/journal.pone.0240769

Chaudhry, A.Z., Lyons, G.E., Gronostajski, R.M., 1997. Expression patterns of the four nuclear factor I genes during mouse embryogenesis indicate a potential role in development. Dev. Dyn. 208, 313–325. 10.1002/(sici)1097-0177(199703)208:3<313::aid-aja3>3.0.co;2-l

Chen, M., Maimaitili, M., Habekost, M., Gill, K.P., Mermet-Joret, N., Nabavi, S., Febbraro, F., Denham, M., 2020. Rapid generation of regionally specified CNS neurons by sequential patterning and conversion of human induced pluripotent stem cells. Stem Cell Res. 48, 101945. 10.1016/j.scr.2020.101945

Ciceri, G., Baggiolini, A., Cho, H.S., Kshirsagar, M., Benito-Kwiecinski, S., Walsh, R.M., Aromolaran, K.A., Gonzalez-Hernandez, A.J., Munguba, H., Koo, S.Y., Xu, N., Sevilla, K.J., Goldstein, P.A., Levitz, J., Leslie, C.S., Koche, R.P., Studer, L., 2024. An epigenetic barrier sets the timing of human neuronal maturation. Nature 626, 881–890. 10.1038/s41586-023-06984-8

Clark, B.S., Stein-O’Brien, G.L., Shiau, F., Cannon, G.H., Davis-Marcisak, E., Sherman, T., Santiago, C.P., Hoang, T.V., Rajaii, F., James-Esposito, R.E., Gronostajski, R.M., Fertig, E.J., Goff, L.A., Blackshaw, S., 2019. Single-Cell RNA-Seq Analysis of Retinal Development Identifies NFI Factors as Regulating Mitotic Exit and Late-Born Cell Specification. Neuron 1–22. 10.1016/j.neuron.2019.04.010

Delás, M.J., Kalaitzis, C.M., Fawzi, T., Demuth, M., Zhang, I., Stuart, H.T., Costantini, E., Ivanovitch, K., Tanaka, E.M., Briscoe, J., 2023. Developmental cell fate choice in neural tube progenitors employs two distinct cis-regulatory strategies. Dev Cell 58, 3–17.e8. 10.1016/j.devcel.2022.11.016

Delás, M.J., Sabin, L.R., Dolzhenko, E., Knott, S.R., Maravilla, E.M., Jackson, B.T., Wild, S.A., Kovacevic, T., Stork, E.M., Zhou, M., Erard, N., Lee, E., Kelley, D.R., Roth, M., Barbosa, I.A., Zuber, J., Rinn, J.L., Smith, A.D., Hannon, G.J., 2017. lncRNA requirements for mouse acute myeloid leukemia and normal differentiation. Elife 6, e25607. 10.7554/elife.25607

Delile, J., Rayon, T., Melchionda, M., Edwards, A., Briscoe, J., Sagner, A., 2019. Single cell transcriptomics reveals spatial and temporal dynamics of gene expression in the developing mouse spinal cord. Development 146, dev173807. 10.1242/dev.173807

Deneen, B., Ho, R., Lukaszewicz, A., Hochstim, C.J., Gronostajski, R.M., Anderson, D.J., 2006. The Transcription Factor NFIA Controls the Onset of Gliogenesis in the Developing Spinal Cord. Neuron 52, 953–968. 10.1016/j.neuron.2006.11.019

Deng, Q., Andersson, E., Hedlund, E., Alekseenko, Z., Coppola, E., Panman, L., Millonig, J.H., Brunet, J.-F., Ericson, J., Perlmann, T., 2011. Specific and integrated roles of Lmx1a, Lmx1b and Phox2a in ventral midbrain development. Development 138, 3399–3408. 10.1242/dev.065482

Denny, S.K., Yang, D., Chuang, C.-H., Brady, J.J., Lim, J.S., Grüner, B.M., Chiou, S.-H., Schep, A.N., Baral, J., Hamard, C., Antoine, M., Wislez, M., Kong, C.S., Connolly, A.J., Park, K.-S., Sage, J., Greenleaf, W.J., Winslow, M.M., 2016. Nfib Promotes Metastasis through a Widespread Increase in Chromatin Accessibility. Cell 1–56. 10.1016/j.cell.2016.05.052

Deska-Gauthier, D., Borowska-Fielding, J., Jones, C., Zhang, H., MacKay, C.S., Michail, R., Bennett, L.A., Bikoff, J.B., Zhang, Y., 2024. Embryonic temporal-spatial delineation of excitatory spinal V3 interneuron diversity. Cell Rep. 43, 113635. 10.1016/j.celrep.2023.113635

Dessaud, E., McMahon, A.P., Briscoe, J., 2008. Pattern formation in the vertebrate neural tube: a sonic hedgehog morphogen-regulated transcriptional network. Development 135, 2489–2503. 10.1242/dev.009324

Doe, C.Q., 2017. Temporal Patterning in the Drosophila CNS. Annu. Rev. Cell Dev. Biol. 33, 219–240. 10.1146/annurev-cellbio-111315-125210

Doetschman, T., Gregg, R.G., Maeda, N., Hooper, M.L., Melton, D.W., Thompson, S., Smithies, O., 1987. Targetted correction of a mutant HPRT gene in mouse embryonic stem cells. Nature 330, 576–578. 10.1038/330576a0

Erclik, T., Li, X., Courgeon, M., Bertet, C., Chen, Z., Baumert, R., Ng, J., Koo, C., Arain, U., Behnia, R., Rodriguez, A.D.V., Senderowicz, L., Negre, N., White, K.P., Desplan, C., 2017. Integration of temporal and spatial patterning generates neural diversity. Nature 541, 365–370. 10.1038/nature20794

Frith, T.J.R., Briscoe, J., Boezio, G.L.M., 2023. From signalling to form: the coordination of neural tube patterning. Curr. Top. Dev. Biol. 10.1016/bs.ctdb.2023.11.004

Fuhrmann, G., Chung, A.C.-K., Jackson, K.J., Hummelke, G., Baniahmad, A., Sutter, J., Sylvester, I., Schöler, H.R., Cooney, A.J., 2001. Mouse Germline Restriction of Oct4 Expression by Germ Cell Nuclear Factor. Dev. Cell 1, 377–387. 10.1016/s1534-5807(01)00038-7

Gao, P., Postiglione, M.P., Krieger, T.G., Hernandez, L., Wang, C., Han, Z., Streicher, C., Papusheva, E., Insolera, R., Chugh, K., Kodish, O., Huang, K., Simons, B.D., Luo, L., Hippenmeyer, S., Shi, S.-H., 2014. Deterministic Progenitor Behavior and Unitary Production of Neurons in the Neocortex. Cell 159, 775–788. 10.1016/j.cell.2014.10.027

Gouti, M., Tsakiridis, A., Wymeersch, F.J., Huang, Y., Kleinjung, J., Wilson, V., Briscoe, J., 2014. In vitro generation of neuromesodermal progenitors reveals distinct roles for wnt signalling in the specification of spinal cord and paraxial mesoderm identity. PLoS Biology 12, e1001937. 10.1371/journal.pbio.1001937

Granja, J.M., Corces, M.R., Pierce, S.E., Bagdatli, S.T., Choudhry, H., Chang, H.Y., Greenleaf, W.J., 2021. ArchR is a scalable software package for integrative single-cell chromatin accessibility analysis. Nature genetics 53, 403–411. 10.1038/s41588-021-00790-6

Gu, P., Xu, X., Menuet, D.L., Chung, A.C. -K., Cooney, A.J., 2011. Differential Recruitment of Methyl CpG-Binding Domain Factors and DNA Methyltransferases by the Orphan Receptor Germ Cell Nuclear Factor Initiates the Repression and Silencing of Oct4. STEM CELLS 29, 1041–1051. 10.1002/stem.652

Gu, Z., Eils, R., Schlesner, M., 2016. Complex heatmaps reveal patterns and correlations in multidimensional genomic data. Bioinformatics 32, 2847–2849. 10.1093/bioinformatics/btw313

Hayashi, M., Hinckley, C.A., Driscoll, S.P., Moore, N.J., Levine, A.J., Hilde, K.L., Sharma, K., Pfaff, S.L., 2018. Graded Arrays of Spinal and Supraspinal V2a Interneuron Subtypes Underlie Forelimb and Hindlimb Motor Control. Neuron 97, 869–884.e5. 10.1016/j.neuron.2018.01.023

Hochstim, C., Deneen, B., Lukaszewicz, A., Zhou, Q., Anderson, D.J., 2008. Identification of Positionally Distinct Astrocyte Subtypes whose Identities Are Specified by a Homeodomain Code. Cell 133, 510–522. 10.1016/j.cell.2008.02.046

Hollyday, M., Hamburger, V., 1977. An autoradiographic study of the formation of the lateral motor column in the chick embryo. Brain Res. 132, 197–208. 10.1016/0006-8993(77)90416-4

Javed, A., Mattar, P., Lu, S., Kruczek, K., Kloc, M., Gonzalez-Cordero, A., Bremner, R., Ali, R.R., Cayouette, M., 2020. Pou2f1 and Pou2f2 cooperate to control the timing of cone photoreceptor production in the developing mouse retina. Development 147, dev188730. 10.1242/dev.188730

Kang, P., Lee, H.K., Glasgow, S.M., Finley, M., Donti, T., Gaber, Z.B., Graham, B.H., Foster, A.E., Novitch, B.G., Gronostajski, R.M., Deneen, B., 2012. Sox9 and NFIA Coordinate a Transcriptional Regulatory Cascade during the Initiation of Gliogenesis. Neuron 74, 79–94. 10.1016/j.neuron.2012.01.024

Kartha, V.K., Duarte, F.M., Hu, Y., Ma, S., Chew, J.G., Lareau, C.A., Earl, A., Burkett, Z.D., Kohlway, A.S., Lebofsky, R., Buenrostro, J.D., 2022. Functional inference of gene regulation using single-cell multi-omics. Cell Genom. 2, 100166. 10.1016/j.xgen.2022.100166

Kohwi, M., Doe, C.Q., 2013. Temporal fate specification and neural progenitor competence during development. Nat. Rev. Neurosci. 14, 823–838. 10.1038/nrn3618

Konstantinides, N., Holguera, I., Rossi, A.M., Escobar, A., Dudragne, L., Chen, Y.-C., Tran, T.N., Jaimes, A.M.M., Özel, M.N., Simon, F., Shao, Z., Tsankova, N.M., Fullard, J.F., Walldorf, U., Roussos, P., Desplan, C., 2022. A complete temporal transcription factor series in the fly visual system. Nature 604, 316–322. 10.1038/s41586-022-04564-w

Kutejova, E., Sasai, N., Shah, A., Gouti, M., Briscoe, J., 2016. Neural Progenitors Adopt Specific Identities by Directly Repressing All Alternative Progenitor Transcriptional Programs. Dev Cell 36, 639–653. 10.1016/j.devcel.2016.02.013

Lim, L., Mi, D., Llorca, A., Marín, O., 2018. Development and Functional Diversification of Cortical Interneurons. Neuron 100, 294–313. 10.1016/j.neuron.2018.10.009

Lin, H.-C., Janssens, J., Kroell, A.-S., Hornauer, P., Santel, M., Okamoto, R., Karava, K., Priouret, M., Garcia, M.P., Schroeter, M., Camp, J.G., Treutlein, B., 2023. Human neuron subtype programming through combinatorial patterning with scRNA-seq readouts. bioRxiv 2023.12.12.571318. 10.1101/2023.12.12.571318

Liu, R., Liu, H., Chen, X., Kirby, M., Brown, P.O., Zhao, K., 2001. Regulation of CSF1 Promoter by the SWI/SNF-like BAF Complex. Cell 106, 309–318. 10.1016/s0092-8674(01)00446-9

Lodato, S., Arlotta, P., 2014. Generating Neuronal Diversity in the Mammalian Cerebral Cortex. Annu. Rev. Cell Dev. Biol. 31, 1–22. 10.1146/annurev-cellbio-100814-125353

Love, M.I., Huber, W., Anders, S., 2014. Moderated estimation of fold change and dispersion for RNA-seq data with DESeq2. Genome biology 15, 550. 10.1186/s13059-014-0550-8

Lu, Q.R., Yuk, D., Alberta, J.A., Zhu, Z., Pawlitzky, I., Chan, J., McMahon, A.P., Stiles, C.D., Rowitch, D.H., 2000. Sonic Hedgehog–Regulated Oligodendrocyte Lineage Genes Encoding bHLH Proteins in the Mammalian Central Nervous System. Neuron 25, 317–329. 10.1016/s0896-6273(00)80897-1

Lucas, T., Hafer, T.L., Zhang, H.G., Molotkova, N., Kohwi, M., 2021. Discrete cis-acting element regulates developmentally timed gene-lamina relocation and neural progenitor competence in vivo. Dev. Cell 56, 2649–2663.e6. 10.1016/j.devcel.2021.08.020

Lyu, P., Hoang, T., Santiago, C.P., Thomas, E.D., Timms, A.E., Appel, H., Gimmen, M., Le, N., Jiang, L., Kim, D.W., Chen, S., Espinoza, D.F., Telger, A.E., Weir, K., Clark, B.S., Cherry, T.J., Qian, J., Blackshaw, S., 2021. Gene regulatory networks controlling temporal patterning, neurogenesis, and cell-fate specification in mammalian retina. Cell Rep. 37, 109994. 10.1016/j.celrep.2021.109994

Manno, G.L., Siletti, K., Furlan, A., Gyllborg, D., Vinsland, E., Albiach, A.M., Langseth, C.M., Khven, I., Lederer, A.R., Dratva, L.M., Johnsson, A., Nilsson, M., Lönnerberg, P., Linnarsson, S., 2021. Molecular architecture of the developing mouse brain. Nature 596, 92–96. 10.1038/s41586-021-03775-x

Mattar, P., Ericson, J., Blackshaw, S., Cayouette, M., 2015. A Conserved Regulatory Logic Controls Temporal Identity in Mouse Neural Progenitors. Neuron 85, 497–504. 10.1016/j.neuron.2014.12.052

Mayer, C., Hafemeister, C., Bandler, R.C., Machold, R., Brito, R.B., Jaglin, X., Allaway, K., Butler, A., Fishell, G., Satija, R., 2018. Developmental diversification of cortical inhibitory interneurons. Nature 555, 457–462. 10.1038/nature25999

Moreau, M.X., Saillour, Y., Cwetsch, A.W., Pierani, A., Causeret, F., 2021. Single-cell transcriptomics of the early developing mouse cerebral cortex disentangles the spatial and temporal components of neuronal fate acquisition. Development 148. 10.1242/dev.197962

Nehme, R., Zuccaro, E., Ghosh, S.D., Li, C., Sherwood, J.L., Pietilainen, O., Barrett, L.E., Limone, F., Worringer, K.A., Kommineni, S., Zang, Y., Cacchiarelli, D., Meissner, A., Adolfsson, R., Haggarty, S., Madison, J., Muller, M., Arlotta, P., Fu, Z., Feng, G., Eggan, K., 2018. Combining NGN2 Programming with Developmental Patterning Generates Human Excitatory Neurons with NMDAR-Mediated Synaptic Transmission. Cell Rep. 23, 2509–2523. 10.1016/j.celrep.2018.04.066

Nishi, Y., Zhang, X., Jeong, J., Peterson, K.A., Vedenko, A., Bulyk, M.L., Hide, W.A., McMahon, A.P., 2015. A direct fate exclusion mechanism by Sonic hedgehog-regulated transcriptional repressors. Development 142, 3286–3293. 10.1242/dev.124636

Novitch, B.G., Chen, A.I., Jessell, T.M., 2001. Coordinate regulation of motor neuron subtype identity and pan-neuronal properties by the bHLH repressor Olig2. Neuron 31, 773–789.

Oosterveen, T., Kurdija, S., Alekseenko, Z., Uhde, C.W., Bergsland, M., Sandberg, M., Andersson, E., Dias, J.M., Muhr, J., Ericson, J., 2012. Mechanistic Differences in the Transcriptional Interpretation of Local and Long-Range Shh Morphogen Signaling. Dev Cell 23, 1006–1019. 10.1016/j.devcel.2012.09.015

OssewardII, P.J., Amin, N.D., Moore, J.D., Temple, B.A., Barriga, B.K., Bachmann, L.C., BeltranJr., F., Gullo, M., Clark, R.C., Driscoll, S.P., Pfaff, S.L., Hayashi, M., 2021. Conserved genetic signatures parcellate cardinal spinal neuron classes into local and projection subsets. Science 372, 385–393. 10.1126/science.abe0690

Peterson, K.A., Nishi, Y., Ma, W., Vedenko, A., Shokri, L., Zhang, X., McFarlane, M., Baizabal, J.-M., Junker, J.P., Oudenaarden, A. van, Mikkelsen, T., Bernstein, B.E., Bailey, T.L., Bulyk, M.L., Wong, W.H., McMahon, A.P., 2012. Neural-specific Sox2 input and differential Gli-binding affinity provide context and positional information in Shh-directed neural patterning. Genes Dev 26, 2802–2816. 10.1101/gad.207142.112

Plachez, C., Lindwall, C., Sunn, N., Piper, M., Moldrich, R.X., Campbell, C.E., Osinski, J.M., Gronostajski, R.M., Richards, L.J., 2008. Nuclear factor I gene expression in the developing forebrain. J. Comp. Neurol. 508, 385–401. 10.1002/cne.21645

Ramírez, F., Ryan, D.P., Grüning, B., Bhardwaj, V., Kilpert, F., Richter, A.S., Heyne, S., Dündar, F., Manke, T., 2016. deepTools2: a next generation web server for deep-sequencing data analysis. Nucleic Acids Res. 44, W160–W165. 10.1093/nar/gkw257

Rowitch, D.H., Kriegstein, A.R., 2010. Developmental genetics of vertebrate glial–cell specification. Nature 468, 214–222. 10.1038/nature09611

Sagner, A., Zhang, I., Watson, T., Lazaro, J., Melchionda, M., Briscoe, J., 2021. A shared transcriptional code orchestrates temporal patterning of the central nervous system. Plos Biol 19, e3001450. 10.1371/journal.pbio.3001450

Sarropoulos, I., Sepp, M., Frömel, R., Leiss, K., Trost, N., Leushkin, E., Okonechnikov, K., Joshi, P., Giere, P., Kutscher, L.M., Cardoso-Moreira, M., Pfister, S.M., Kaessmann, H., 2021. Developmental and evolutionary dynamics of cis-regulatory elements in mouse cerebellar cells. Science 373. 10.1126/science.abg4696

Sato, N., Kondo, M., Arai, K., 2006. The orphan nuclear receptor GCNF recruits DNA methyltransferase for Oct-3/4 silencing. Biochem. Biophys. Res. Commun. 344, 845–851. 10.1016/j.bbrc.2006.04.007

Scott, C.E., Wynn, S.L., Sesay, A., Cruz, C., Cheung, M., Gaviro, M.-V.G., Booth, S., Gao, B., Cheah, K.S.E., Lovell-Badge, R., Briscoe, J., 2010. SOX9 induces and maintains neural stem cells. Nat. Neurosci. 13, 1181–1189. 10.1038/nn.2646

Sen, S.Q., 2023. Generating neural diversity through spatial and temporal patterning. Semin. Cell Dev. Biol. 142, 54–66. 10.1016/j.semcdb.2022.06.002

Sen, S.Q., Chanchani, S., Southall, T.D., Doe, C.Q., 2019. Neuroblast-specific open chromatin allows the temporal transcription factor, Hunchback, to bind neuroblast-specific loci. eLife 8, e44036. 10.7554/elife.44036

Shu, M., Hong, D., Lin, H., Zhang, J., Luo, Z., Du, Y., Sun, Z., Yin, M., Yin, Y., Liu, L., Bao, S., Liu, Z., Lu, F., Huang, J., Dai, J., 2022. Single-cell chromatin accessibility identifies enhancer networks driving gene expression during spinal cord development in mouse. Dev. Cell 57, 2761–2775.e6. 10.1016/j.devcel.2022.11.011

Sockanathan, S., Jessell, T.M., 1998. Motor Neuron–Derived Retinoid Signaling Specifies the Subtype Identity of Spinal Motor Neurons. Cell 94, 503–514. 10.1016/s0092-8674(00)81591-3

Stolt, C.C., Lommes, P., Sock, E., Chaboissier, M.-C., Schedl, A., Wegner, M., 2003. The Sox9 transcription factor determines glial fate choice in the developing spinal cord. Genes Dev. 17, 1677–1689. 10.1101/gad.259003

Sun, K., Liu, X., Lan, X., 2024. A single-cell atlas of chromatin accessibility in mouse organogenesis. Nat. Cell Biol. 26, 1200–1211. 10.1038/s41556-024-01435-6

Tchieu, J., Calder, E.L., Guttikonda, S.R., Gutzwiller, E.M., Aromolaran, K.A., Steinbeck, J.A., Goldstein, P.A., Studer, L., 2019. NFIA is a gliogenic switch enabling rapid derivation of functional human astrocytes from pluripotent stem cells. Nat. Biotechnol. 37, 267–275. 10.1038/s41587-019-0035-0

Telley, L., Agirman, G., Prados, J., Amberg, N., Fièvre, S., Oberst, P., Bartolini, G., Vitali, I., Cadilhac, C., Hippenmeyer, S., Nguyen, L., Dayer, A., Jabaudon, D., 2019. Temporal patterning of apical progenitors and their daughter neurons in the developing neocortex. Science 364. 10.1126/science.aav2522

Tiberi, L., Vanderhaeghen, P., Ameele, J. van den, 2012. Cortical neurogenesis and morphogens: diversity of cues, sources and functions. Curr. Opin. Cell Biol. 24, 269–276. 10.1016/j.ceb.2012.01.010

Tiklová, K., Björklund, Å.K., Lahti, L., Fiorenzano, A., Nolbrant, S., Gillberg, L., Volakakis, N., Yokota, C., Hilscher, M.M., Hauling, T., Holmström, F., Joodmardi, E., Nilsson, M., Parmar, M., Perlmann, T., 2019. Single-cell RNA sequencing reveals midbrain dopamine neuron diversity emerging during mouse brain development. Nat. Commun. 10, 581. 10.1038/s41467-019-08453-1

Vierbuchen, T., Ling, E., Cowley, C.J., Couch, C.H., Wang, X., Harmin, D.A., Roberts, C.W.M., Greenberg, M.E., 2017. AP-1 Transcription Factors and the BAF Complex Mediate Signal-Dependent Enhancer Selection. Mol. Cell 68, 1067–1082.e12. 10.1016/j.molcel.2017.11.026

Vierstra, J., Lazar, J., Sandstrom, R., Halow, J., Lee, K., Bates, D., Diegel, M., Dunn, D., Neri, F., Haugen, E., Rynes, E., Reynolds, A., Nelson, J., Johnson, A., Frerker, M., Buckley, M., Kaul, R., Meuleman, W., Stamatoyannopoulos, J.A., 2020. Global reference mapping of human transcription factor footprints. Nature 583, 729–736. 10.1038/s41586-020-2528-x

Worthy, A.E., Anderson, J.T., Lane, A.R., Gomez-Perez, L., Wang, A.A., Griffith, R.W., Rivard, A.F., Bikoff, J.B., Alvarez, F.J., 2023. SPINAL V1 INHIBITORY INTERNEURON CLADES DIFFER IN BIRTHDATE, PROJECTIONS TO MOTONEURONS AND HETEROGENEITY. bioRxiv 2023.11.29.569270. 10.1101/2023.11.29.569270

Xu, C., Ramos, T.B., Marshall, O.J., Doe, C.Q., 2024. Notch signaling and Bsh homeodomain activity are integrated to diversify Drosophila lamina neuron types. eLife 12, RP90136. 10.7554/elife.90136

Xu, H.-T., Han, Z., Gao, P., He, S., Li, Z., Shi, W., Kodish, O., Shao, W., Brown, K.N., Huang, K., Shi, S.-H., 2014. Distinct Lineage-Dependent Structural and Functional Organization of the Hippocampus. Cell 157, 1552–1564. 10.1016/j.cell.2014.03.067

Yang, Y., Gomez, N., Infarinato, N., Adam, R.C., Sribour, M., Baek, I., Laurin, M., Fuchs, E., 2023. The pioneer factor SOX9 competes for epigenetic factors to switch stem cell fates. Nat. Cell Biol. 25, 1185–1195. 10.1038/s41556-023-01184-y

Yu, G., Wang, L.-G., Han, Y., He, Q.-Y., 2012. clusterProfiler: an R Package for Comparing Biological Themes Among Gene Clusters. OMICS: A J. Integr. Biol. 16, 284–287. 10.1089/omi.2011.0118

Yuan, W., Ma, S., Brown, J.R., Kim, K., Murek, V., Trastulla, L., Meissner, A., Lodato, S., Shetty, A.S., Levin, J.Z., Buenrostro, J.D., Ziller, M.J., Arlotta, P., 2022. Temporally divergent regulatory mechanisms govern neuronal diversification and maturation in the mouse and marmoset neocortex. Nat. Neurosci. 25, 1049–1058. 10.1038/s41593-022-01123-4

Zhou, Q., Wang, S., Anderson, D.J., 2000. Identification of a Novel Family of Oligodendrocyte Lineage-Specific Basic Helix–Loop–Helix Transcription Factors. Neuron 25, 331–343. 10.1016/s0896-6273(00)80898-3

